# RAB12-LRRK2 Complex Suppresses Primary Ciliogenesis and Regulates Centrosome Homeostasis in Astrocytes

**DOI:** 10.1101/2024.07.17.603999

**Authors:** Xingjian Li, Hanwen Zhu, Bik Tzu Huang, Xianting Li, Heesoo Kim, Haiyan Tan, Yuanxi Zhang, Insup Choi, Junmin Peng, Pingyi Xu, Ji Sun, Zhenyu Yue

## Abstract

Leucine-rich repeat kinase 2 (LRRK2) phosphorylates a subset of RAB GTPases, and the phosphorylation levels are elevated by Parkinson’s disease (PD)-linked mutations of LRRK2. However, the precise function of the specific RAB GTPase targeted by LRRK2 signaling in the brain remains to be elucidated. Here, we identify RAB12 as a robust LRRK2 substrate in the mouse brains through phosphoproteomics profiling and solve the structure of RAB12-LRRK2 protein complex through Cryo-EM analysis. Mechanistically, RAB12 cooperates with LRRK2 to inhibit primary ciliogenesis and regulate centrosome homeostasis in astrocytes through enhancing the phosphorylation of RAB10 and recruiting Rab interacting lysosomal protein like 1 (RILPL1), while the functions of RAB12 require a direct interaction with LRRK2 and LRRK2 kinase activity. Furthermore, the ciliary deficits and centrosome alteration caused by the PD-linked LRRK2-G2019S mutation are prevented by the deletion of *Rab12* in astrocytes. Thus, our study reveals a physiological function of the RAB12-LRRK2 complex in regulating ciliogenesis and centrosome homeostasis. The RAB12-LRRK2 structure offers a guidance in the therapeutic development of PD by targeting the RAB12-LRRK2 interaction.

## Introduction

The leucine rich repeat kinase 2 (LRRK2) variants cause the most common inherited forms of Parkinson’s disease (PD)^1, 2^. *Lrrk2* encodes a large, complex protein containing both GTPase and kinase domains^3^. High-resolution cryo-EM structures of full-length LRRK2 protein were determined, and the kinase domain was shown to adopt an inactive conformation^4, 5^, suggesting a highly regulated kinase activity. Multiple pathogenic variants of LRRK2 were identified and shown to enhance LRRK2 kinase activity in vitro^6^. Among those disease mutants, the LRRK2-G2019S is the most prevalent^7, 8, 9^. Available evidence has shown that LRRK2 regulates multiple cellular pathways, including immune response, vesicle trafficking, lysosome homeostasis, and synaptic transmission^3, 10, 11^.

Previous proteomics analysis reported that LRRK2 phosphorylates a subset of RAB GTPase at a conserved residue threonine or serine in their switch-II domains, resulting in a potential inhibition of RAB activation and the effector protein binding^12, 13^. The phospho-RAB has a profound impact on their functions as evidenced in impaired autophagosome motility in the axons^14^, primary ciliogenesis^15, 16, 17, 18, 19, 20, 21, 22^, and centriolar cohesion^19, 23^. For example, LRRK2 was shown to regulate primary ciliogenesis through RAB8 and RAB10 and their interactions with effector Rab interacting lysosomal protein like 1/2 (RILPL1/2)^16, 18, 24^. Subsequent studies verified that LRRK2 pathogenic variants R1441C and G2019S interfere with primary cilia formation associated with an impairment of sonic hedgehog signaling^15^.

The identification of the selective RAB GTPase as the targets of LRRK2 kinase began to provide important insight into the molecular mechanism underlying LRRK2 pathogenic signaling pathways^10, 25,26^. However, the exact mechanism for how multiple RAB proteins participate in LRRK2 pathogenic pathways remains to be clarified. Interestingly, study of *Lrrk2*-G2019S knockin (KI) mouse tissues demonstrated an increase of phospho-RAB12 levels, while the change of phosphorylation levels of other RAB proteins is not understood^27^, implicating RAB12 as an important substrate of LRRK2 in the brain at basal condition. RAB12 can be recruited to LRRK2-associated lysosomes or endolysosomes and are phosphorylated by LRRK2^28, 29, 30^. Two most recent reports showed that RAB12 is an activator of LRRK2 kinase activity that can promote lysosome recruitment of LRRK2 and facilitate LRRK2 phosphorylation of RAB10 in cell cultures from non-brain cell types^22, 31^. Despite the notions, the physiological function of RAB12 and precise functional relationship between LRRK2, RAB12, and other RAB GTPase remains to be elucidated particularly in neurons and glia.

Here we report RAB12 as a physiological substrate of LRRK2 in the brain, reveal the cryo-EM structure of RAB12-LRRK2 protein complex, and characterize the function of RAB12 in the glia. We show that RAB12 is an inhibitor of primary ciliogenesis, which requires a direct binding with LRRK2. Our study provides an insight into the potential mechanism whereby RAB12-LRRK2 complex controls the homeostasis of astrocytic ciliogenesis and centrosomes. Furthermore, our study shows the role of RAB12 in mediating the pathogenic function of LRRK2-G2019S in astrocytes. Our study thus establishes the physiological function of the RAB12-LRRK2 complex and provides a structural framework for targeting RAB12-LRRK2 interaction in therapeutic development.

## Results

### 1. Structure of the RAB12–LRRK2 complex

To identify the physiological substrates of LRRK2 kinase in the brain, we performed a deep proteome and phosphoproteomics analysis using brain lysates from *Lrrk2* knockout (KO), *Lrrk2* BAC transgenic mice overexpressing *Lrrk*2 wildtype (WTtg), PD-mutant G2019S (GStg)^32^, and control mice (WT). Through the TMT-LC/LC-MS/MS analysis, we identified a total of 10,195 proteins and 72,766 phosphopeptides (52,187 phosphorylation sites out of 7,829 phosphoproteins) (**Fig. S1a-c, Tables. 1-3**). The data revealed that the relative phosphorylation levels at multiple sites on LRRK2 (S910, S935, S955, and S973), OXR1 (S15, S308, and S396), and RAB12 (S105) were correlated with the LRRK2 levels/activities (**Fig. S1c**). We validated the RAB12 phosphorylation levels in the above mutant mice through immunoblot analysis (**Fig. S1d**). The above observation suggested that RAB12 is a robust substrate of LRRK2 in the brain under basal condition.

To characterize the interaction between RAB12 and LRRK2 at atomic details, we solved the cryo-electron microscopy (cryo-EM) structures of the human RAB12–LRRK2 complex (**Fig. S2 and Table. S4**). We introduced the GTPase-deficient mutation (active form), Q101L^33^, to RAB12 to obtain homogenous and stable RAB12-LRRK2 complexes for structural analysis. Like LRRK2-alone structures^4^, the RAB12– LRRK2 complex was also captured in both monomer and dimer states at overall resolutions of 4.1 Å and 3.9 Å, respectively (**Fig. S2a),** and LRRK2 protomers in both states adopt an inactive conformation as previously reported^4, 34^ (**Fig. S3a**). Focused refinement was performed for the RAB12-LRRK2 interface to improve the local resolution (**Fig. S2a**). The GTP analog-bound RAB12 resembles RAB29 and RAB32 in their complexes with LRRK2 (**Fig. 1b and Fig. S3b**).

**Fig. 1.**
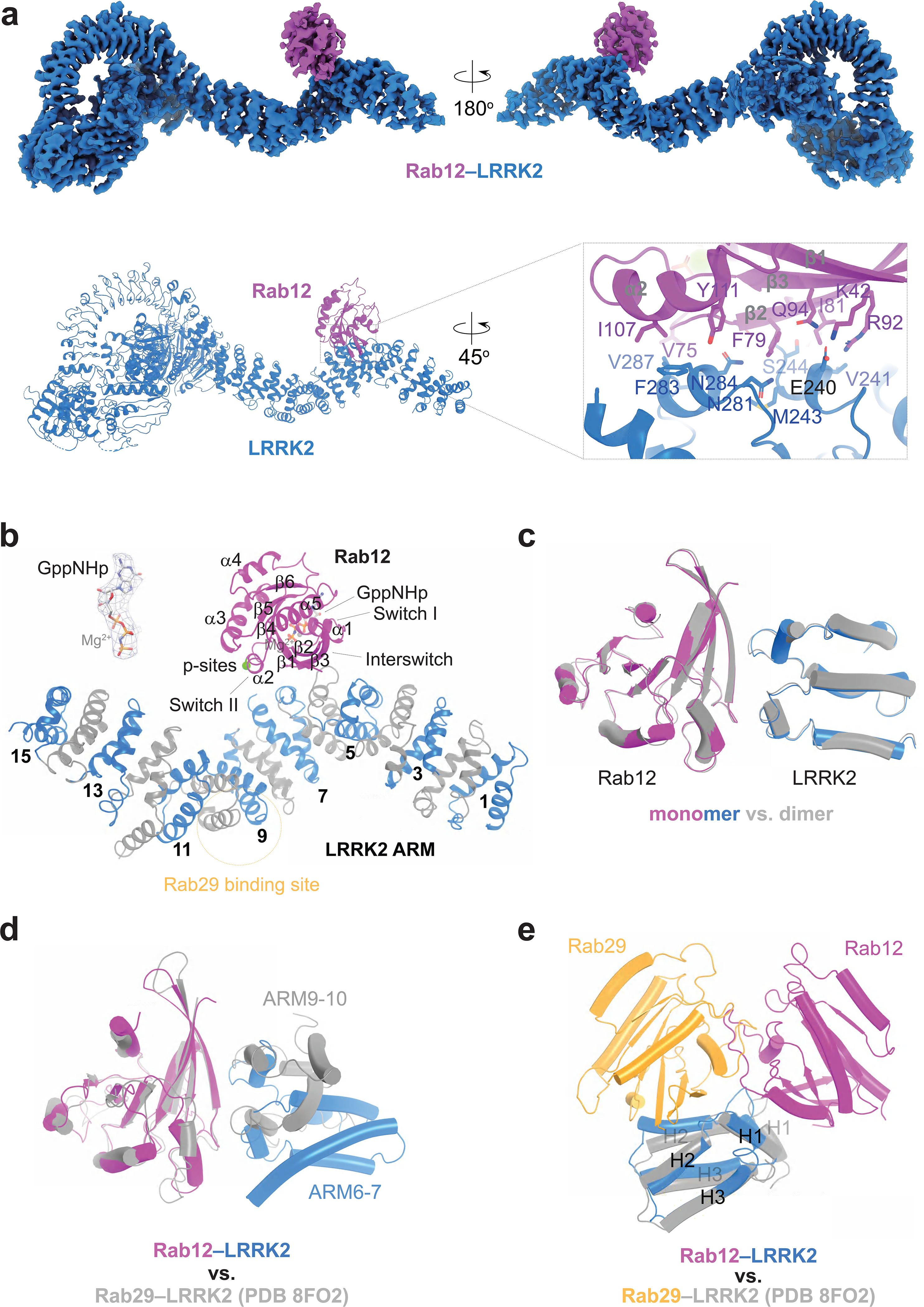
**Structural analysis of the RAB12–LRRK2 complex. a**, The cryo-EM map and atomic model of the RAB12–LRRK2 complex. RAB12 and LRRK2 are colored in purple and blue, respectively. Side chains involved in interactions between RAB12 and LRRK2 are shown as sticks. **b,** Interactions between the LRRK2 ARM domain and RAB12. The ARM repeats of LRRK2 are indicated by numbers. Key structural elements in RAB12 are labeled. The phosphorylation site (p-site) is shown as green sphere. Cryo-EM density of the GTP analog and Mg2+ bound to RAB12 is shown. **c,** Structural comparison of the RAB12–LRRK2 interfaces between RAB12-LRRK2 monomer and dimer. **d,** Superposition analysis of the RAB12–LRRK2 and RAB29–LRRK2 (PDB 8FO2) interfaces based on RAB proteins. **e,** Superposition analysis of the RAB12–LRRK2 and RAB29–LRRK2 (PDB 8FO2) interfaces based on the LRRK2 ARM repeats. The three helices (H1, H2, and H3) of each ARM repeat are labeled.

The interaction interfaces between RAB12 and LRRK2 in both monomers and dimers are almost identical and mediated by CDR1 and Switch I-Interswitch-Switch II motifs of RAB12 and ARM6-ARM7 of LRRK2 (**Fig. 1a-c**). Key interface residues are annotated in **Fig. 1a**. Comparing the RAB12–LRRK2 and RAB29/RAB32–LRRK2 complexes, we found that all RABs use the CDR1 and Switch I-Interswitch-Switch II motifs for LRRK2 binding^34^, though the α2 helix in Rab12 seems to make closer contacts with ARM repeats (**Fig. 1d and Fig. S3c-d**). As a result, Tyr111 of RAB12 sits in the middle of the interface, while the side chain of corresponding residue Tyr77 points away from the ARM domain in RAB29 (**Fig. S3c-d**). On the other hand, RAB12 binds to the H1 helices of ARM6-ARM7 while RAB29 and RAB32 mainly interact with the H2 helices of ARM9-ARM10 of LRRK2 (**Fig. 1e and S3f**). Of note, each ARM repeat contains three helices: H1-H3. Sequence alignment revealed that only a few interface residues are not conserved among the three RAB GTPases, including Lys42, Ile81 and Ile107 of RAB12 and Leu76 of RAB29, resulting in a totally different binding surface on the ARM domain of LRRK2 (**Fig. S3c- d, e-f**). Glu240 of LRRK2 interacts with two of the three RAB12-specific residues (Lys42 and Ile81), forms salt bridges with Arg92 and Lys42 of RAB12 (**Fig. S3c**) and should be critical for RAB12-LRRK2 interactions.

#### Lack of *Rab12* increases the number of ciliated astrocytes in vivo and in vitro

To elucidate the physiological function of *Rab12* in the brain, we generated *Rab12* KO mice by deleting the exon 3 of *Rab12* from the genomic sequence using the CRISPR-Cas9 gene editing (**Fig. S4a**). The *Rab12* KO mice are viable and show no apparent developmental defect (not shown). Given the evidence for the involvement of LRRK2 in primary ciliogenesis^13, 15, 16, 17, 18, 20, 24^, we tested the effects of *Rab12* deletion on the primary cilia formation in mouse brains. Staining of *Rab12* KO brain slices with anti-Arl13b antibody showed a significant increase of the ciliated cell percentage in the striatum and cortex and a trend of increase of cilia length in the striatum. However, the average cilia volume of the mutant mice changed little in both regions (**Fig. 2a**). To ascertain the cell types affected in *Rab12* KO brains, we performed co-staining with anti-Arl13b and the antibodies specific for astrocyte or neuron, two main ciliated cell types in the brains^35^. The results indicated that the astrocytes had higher ciliated cell percentage yet no significant change in the cilia length or volume in the mutant brains (striatum and cortex) compared to the controls (**Fig. 2b**). In contrast, the neurons from *Rab12* KO brains showed no difference in the ciliated cell percentage but increased cilia length and volume in the striatum, not in the cortex, compared to the control mice (**Fig. S4b**). Consistently, primary astrocyte cultures derived from *Rab12* KO brains also showed elevated percentage of ciliation but no change of the cilia length and volume (**Fig. 2c**). Therefore, our in vivo and in vitro data indicate that loss of *Rab12* results in an increased number of ciliated astrocytes.

**Fig. 2.**
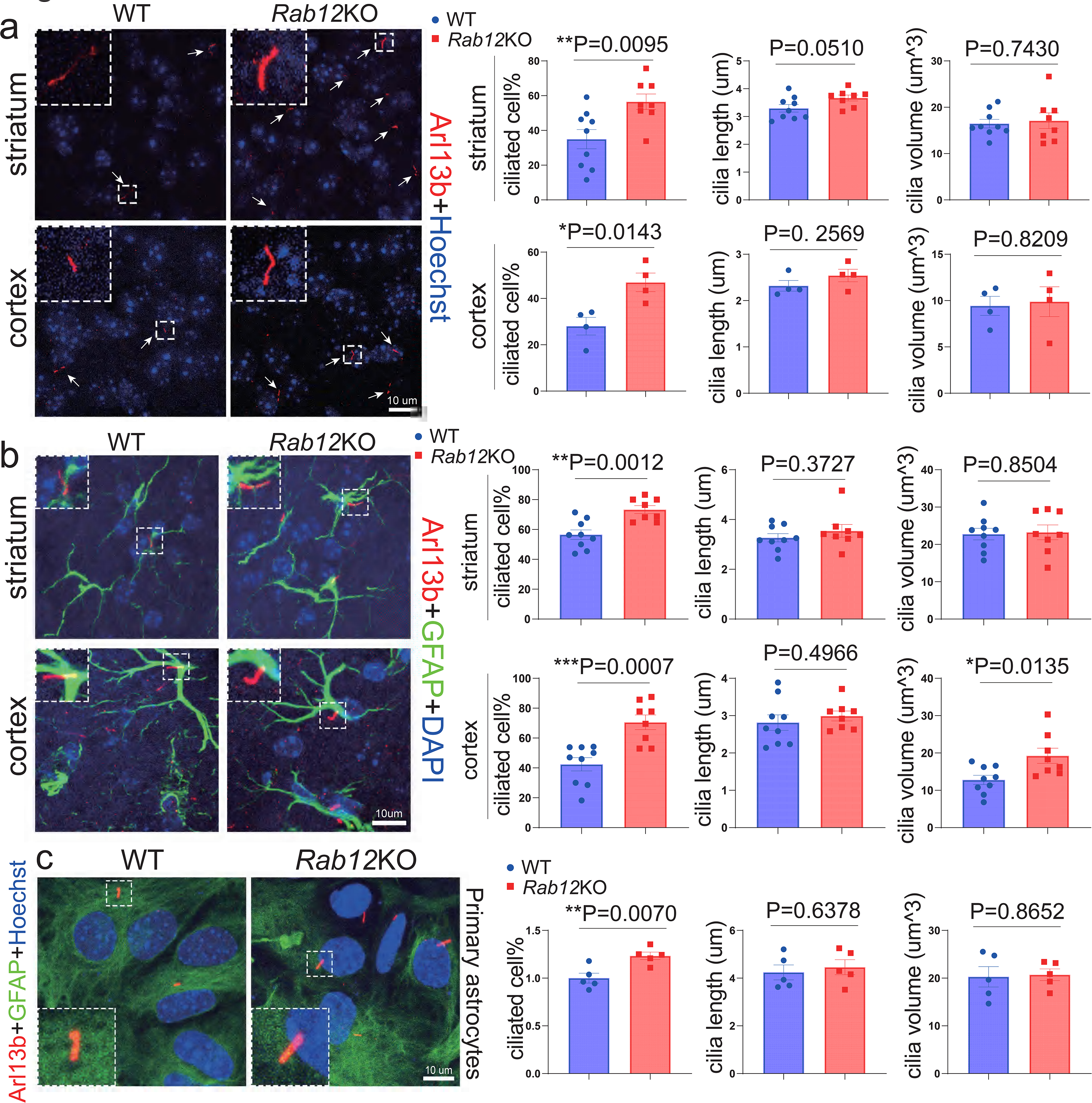
**Characterization of ciliated cells and cilium size *in vivo* and *in vitro*. a**, The primary cilia were detected with anti-Arl13b antibody (white arrowheads) and imaged in the striatum and frontal cortex from wildtype (WT) and *Rab12* KO mice. The ciliated cell percentage, cilia length, and cilia volume were quantified. The boxed images of cilia were enlarged at the upper left corners in each panel. **b**, The primary cilia were detected in the astrocytes (anti-GFAP) in the striatum and frontal cortex of WT and *Rab12* KO mice, and ciliated astrocyte percentage and cilium size were quantified. The boxed images of cilia were enlarged at the upper left corners in each panel. **c**, The cilia in primary cultured astrocytes from neonatal *Rab12* KO mice and littermate controls were examined as in (**b**). The boxed images of cilia were enlarged at the bottom left corners in each panel. Each dot in the graphs represents an individual mouse (**a-b**) and > 60 cells from each mouse (**c**). *P* values were calculated using unpaired two-tailed *t*-tests; \**P*<0.05, \*\**P*<0.01, \*\*\**P*<0.001; error bars = SEM; scale bars are indicated.

#### *Rab12* regulates the homeostasis of primary cilia and centrosomes

We next tested the idea that RAB12 suppresses primary ciliogenesis by focusing on astrocytes. We overexpressed *Rab12* gene in astrocytes by AAV-mediated delivery of HA-tagged *Rab12* into primary cultures. HA-RAB12 overexpression often resulted in large-size amorphic structures with short “spikes” protruding from the surface with anti-Arl13b staining in contrast to GFP-overexpressed astrocytes (**Fig. 3a**). The average intensity of Arl13b fluorescence of the spike-like structures in HA-RAB12- overexpressed astrocytes, however, is less than the cilia (Arl13b+) in GFP-overexpressed cells (**Fig. 3a**). Staining with anti-γ-tubulin, a marker for centrosome which forms basal bodies and sequentially templates cilia formation^36^, showed a significant overlap of γ-tubulin with Arl13b and HA-RAB12 in the large structures. The above staining pattern in HA-RAB12 overexpressed astrocytes is in stark contrast with non-infected or *Gfp*-infected astrocytes, which showed typical primary cilia linked to centrosomes with both γ-tubulin and Arl13b staining (**Fig. 3a**). Indeed, HA-RAB12 overexpression led to a significant increase of γ-tubulin staining volume, compared to GFP overexpression. Using another centrosome specific marker anti-Pericentrin antibody we validated the greater volume of centrosomes (Pericentrin+/HA-RAB12+) in the HA-RAB12 overexpressed astrocytes (**Fig. 3b**). The observation suggests an expansion of centrosomes and perhaps aberrant cilia formation resulted from the overexpression of RAB12.

**Fig. 3.**
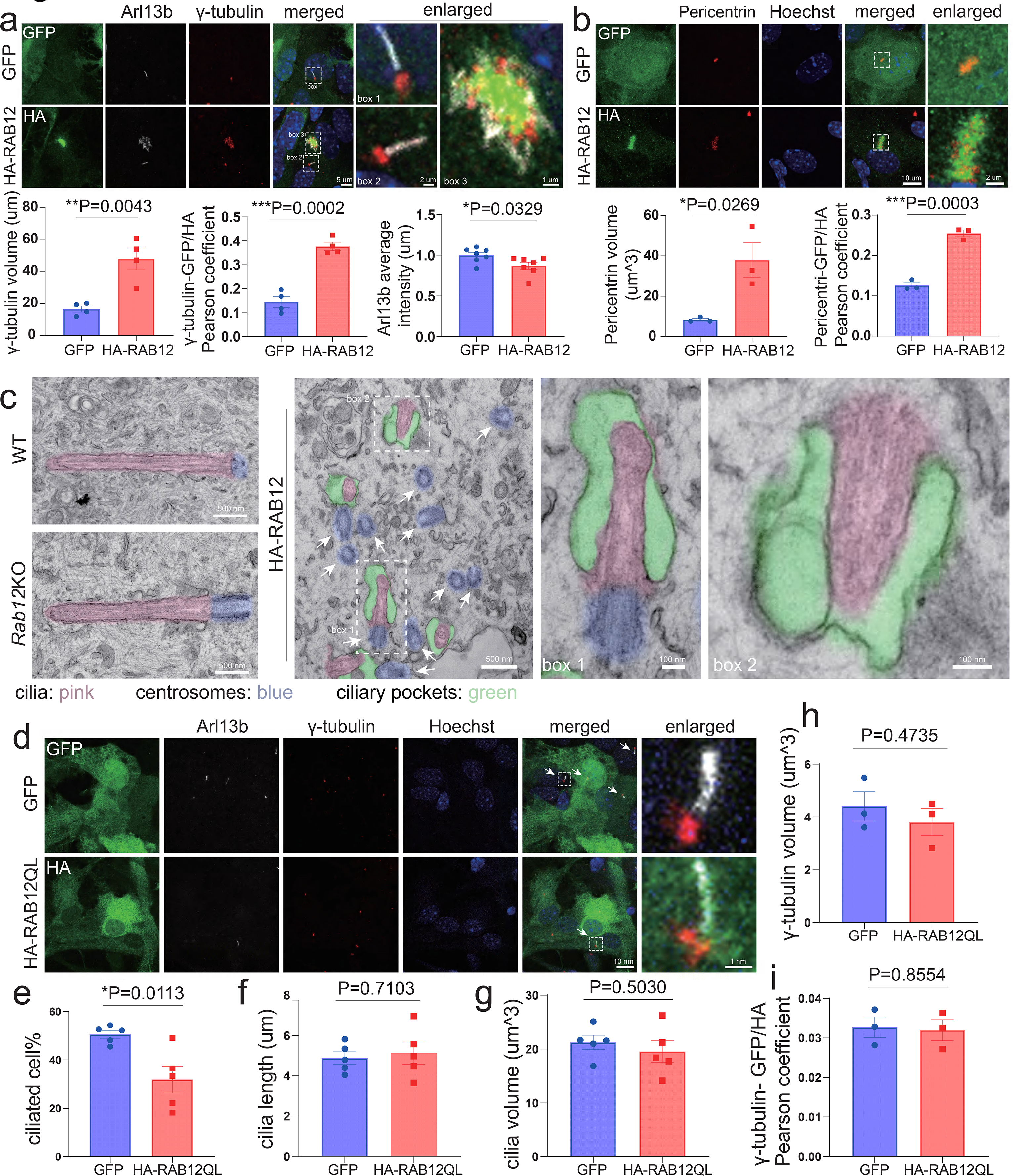
The effect of RAB12 overexpression on primary cilia and homeostasis of centrosomes in astrocytes. Primary culture of astrocytes was infected with AAVs harboring GFAP promoters to express GFP, RAB12WT (WT_OE, **a**-**c**), or the active RAB12 Q100L variant (QL_OE, **d**) conjugated with an HA tag. **a-b**, The astrocytes overexpressing HA-RAB12WT or *Gfp* were marked with HA or GFP antibodies by IF, accordingly. Their primary cilia and centrosomes were detected using Arl13b antibody and γ-tubulin antibody, respectively. The regions of cells containing centrosomes and typical/failed cilia are marked by white-dash boxes and enlarged (**a**-**b**). The graphs at the bottom of (**a**-**b**) are the quantification of γ-tubulin volume, γ-tubulin–GFP/HA Pearson coefficient, and Arl13b fluorescent intensity from the images of (**a**), and Pericentrin volume, and Pericentrin–GFP/HA Pearson coefficient value from images of (**b**). **c**, Electron microscope images from WT, *Rab12* KO, and HA-RAB12WT (WT_OE) overexpressed astrocytes. Centrosomes are indicated in blue, cilia are in pink, and ciliary pockets are in green. The arrowheads indicate the amplificated centrosomes, and the representatives of failed cilia in box 1 and 2 are enlarged and displayed in the right panels. **d**, The cilia and centrosomes in the astrocytes overexpressing GFP*, or* RAB12QL were detected by IF, and the representative cilia from each group were enlarged. **e**-**i**, The qualification of ciliated cell percentage, cilia length, cilia volume, γ-tubulin volume, and γ-tubulin – GFP/RAB12 Pearson coefficient from images as in (**d**). Each dot in the graphs represents >60 cells from each mouse. Unpaired two-tailed *t*-tests, \**P*<0.05, \*\**P*<0.01, \*\*\**P*<0.001; error bars = SEM; scale bars are indicated.

To investigate the details of the cilia alterations, we performed electron microscopy (EM) analysis of the infected astrocytes. In control and *Rab12* KO astrocytes, the normal primary cilia are associated with the centrosome at base (**Fig. 3c**). In contrast, we found frequent clustering of centrosomes/centrioles in HA- *Rab12* infected cells, in which the “dwarf” cilia are associated with aberrant cilia membranes and enlarged lumen enwrapping the abnormal axoneme (**Fig. 3c**). The clustering of centrosomes/centrioles from the EM analysis is consistent with the large volume of γ-tubulin/Pericentrin labeled structures under the confocal microscopy (**Fig. 3a**). They are analogous to what was previously described as centrosome amplification or supernumerary centrioles associated with tumor cells^37, 38^. Thus, our EM result suggested the failure of primary cilia formation in HA*-*RAB12 overexpressed astrocytes. Our observation agreed with a previous report, which suggested that centrosome amplification can inhibit cilia formation or cause clustered, diluted cilia (e.g. reduced Arl13b levels) ^37^.

Moreover, overexpression of HA-RAB12-Q100L (HA-RAB12QL) mouse mutant, a constitutively active form of RAB12^39, 40^, in astrocytes markedly reduced the ciliated cell percentage (**Fig. 3d-g**). However, in contrast to the wildtype, the active RAB12 had little effect on the γ-tubulin staining volume (**Fig. 3h**) and showed no significant co-localization with γ-tubulin staining (**Fig. 3i**). The inhibitory effect of active RAB12 variant on ciliated number of astrocytes corroborates the findings with RAB12 wildtype (WT) overexpression, which stunted cilia formation (**Fig. 3a-c**), while the active RAB12 variant appears not to cause centrosome amplification or associate with centrosomes.

We next asked if overexpression of LRRK2 or other LRRK2 substrates, such as RAB10 and RAB8a, affects the homeostasis of primary cilia and centrosomes in astrocytes. Interestingly, overexpression of RAB10, RAB8a, or LRRK2 influenced neither cilia morphology nor the volume of centrosomes (F**ig. S5a- b**). However, overexpression of RILPL1, an effector of RAB proteins, caused increased centrosome volume (**Fig. S5b**). Therefore, the effect of overexpression on primary cilia and centrosome homeostasis appears to be specific for RAB12, which may act through the effector RILPL1.

#### The RAB12-LRRK2 complex regulates primary cilia and centrosomes

Our structural analysis showed RAB12-LRRK2 complex formation and the binding interface between the two proteins (**Fig. 1**). Therefore, we next tested the role of RAB12-LRRK2 complex in regulating primary cilia and centrosomes in astrocytes, and we postulated that mutations of the critical residues of the interface in RAB12 disrupt the RAB12-LRRK2 interaction and consequently abolish the effects in suppressing ciliogenesis or promoting centrosome amplification. The analysis suggested that K42, F79, R92, and Y111 positioning at the interface of RAB12 are important for binding the ARM of LRRK2 (**Fig. 1a**). To this end, we engineered K42A and Y111A in the active RAB12 variant (HA-RAB12QL) as well as WT (HA-RAB12). Firstly, we verified that both active and WT RAB12 carrying K42A or Y111A lost the interactions with LRRK2 through co-immunoprecipitation (Co-IP) assay (**Fig. 4a-b**). Secondly, in contrast to HA-RAB12QL, overexpression of double mutants HA-RAB12QL-K42A or HA-RAB12QL-Y111A in astrocytes had no impact on the ciliated cell percentage. They did not influence cilia length or cilia volume (**Fig. 4c**). Thirdly, overexpression of HA-RAB12QL in *Lrrk2* KO astrocytes appeared to alleviate the effect of reduced ciliated cell percentage as observed in WT astrocytes, while no influence was found in cilia length or volume (**Fig. 4d, Fig. S6b**). Thus, the role of RAB12 in the suppression of primary ciliogenesis in astrocytes depends on the presence of LRRK2 and requires RAB12-LRRK2 interaction.

**Fig. 4.**
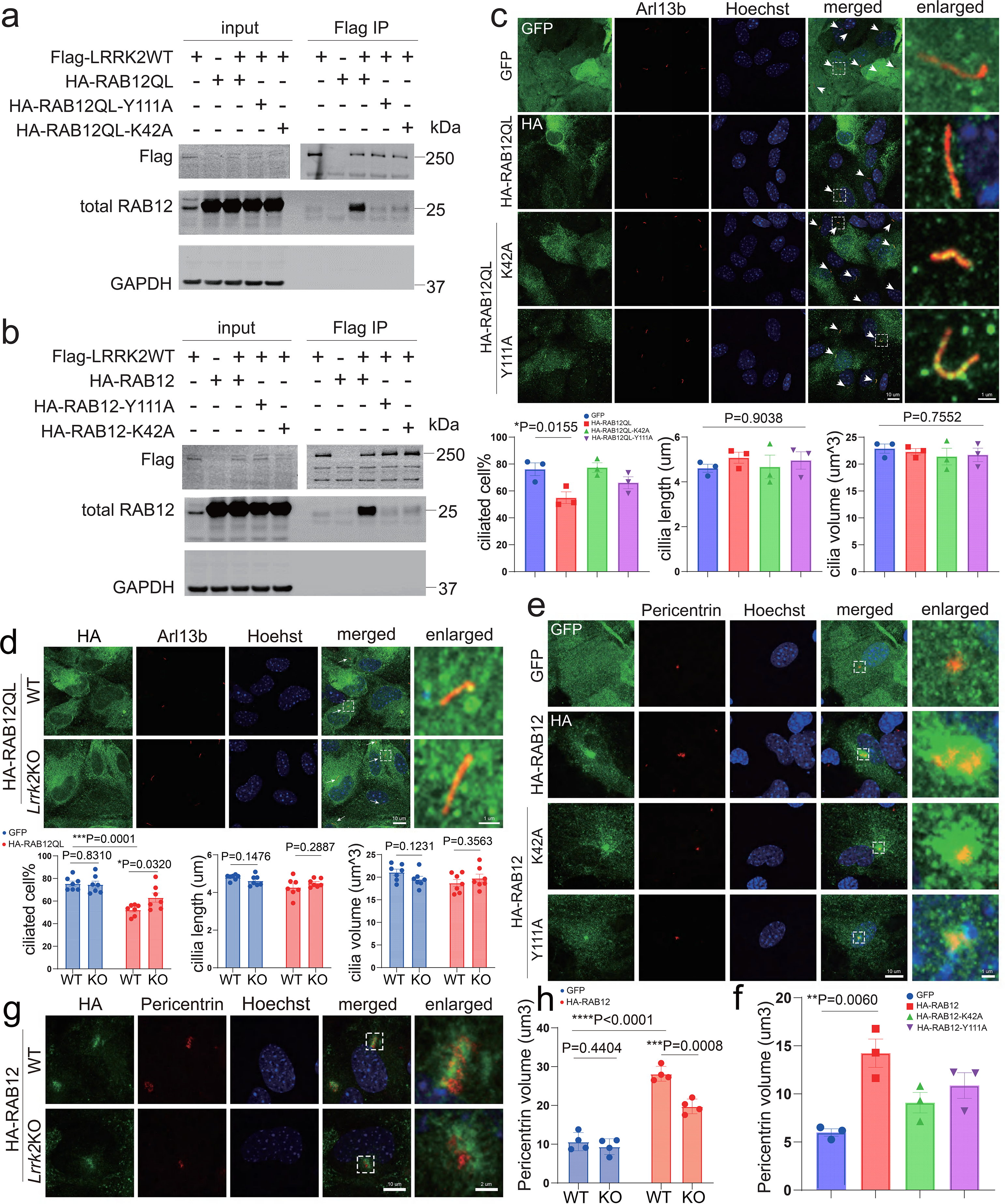
**Examination of the effect of RAB12-LRRK2 binding on cilia formation and centrosome homeostasis. a-b**, The human *LRRK2*WT plasmids were co-transfected with the human *RAB12* variant plasmids, such as HA-*RAB12*QL-K42A, HA-*RAB12*QL-Y111A, HA-*RAB12*WT K42A, or HA- *RAB12*WT Y111A into HEK293T cells. The interactions between LRRK2 and RAB12 or its mutants were examined by co-immunoprecipitation (Co-IP) assay. **a,** The Co-IP between LRRK2 and RAB12QL or its binding deficient mutants. **b,** The Co-IP between LRRK2 and RAB12WT or its binding deficient mutants. **c**, *Gfp*, HA-*RAB12*QL, HA-*RAB*12QL-K42A, or HA-*RAB*12QL-Y111A were transduced into astrocytes by AAVs carrying *GFAP* promotors. **d**, WT and *Lrrk2* KO astrocytes were infected with AAVs carrying HA-*Rab12*QL. The cilia of the infected astrocytes from (**c**-**d**) were detected and their ciliated cell percentage, cilia length, and cilia volume were measured (**c-d**). **e-f**, Primary astrocytes were infected with AAV-*GFAP*-*Gfp*, AAV-*GFAP*-HA-*RAB12*WT, AAV-*GFAP*-HA-*RAB*12WT-K42A, or AAV-*GFAP*-HA-*RAB12*WT-Y111A. The infected astrocytes’ centrosome volume was determined. **g-h**, The centrosome volume in WT and *Lrrk2* KO astrocytes infected with AAV-*GFAP*-*Gfp*, or AAV-*GFAP*-HA- *Rab12*WT was measured. The representative cilia from (**c-d**) and centrosomes from (**e, g**) were enlarged in each panel. Each dot in the graphs represents >60 cells from each mouse. *P* values were calculated by one-way ANOVA (**c, f**) and unpaired two-tailed *t*-tests (**d, h**). Turkey’s HSD post hoc tests were performed when *P* values from one-way ANOVA < 0.05. \**P*<0.05, \*\**P*<0.01, \*\*\**P*<0.001, \*\*\*\**P*<0.0001; error bars = SEM; scale bars are indicated.

We next asked whether LRRK2-binding mutants of RAB12 are affected in the centrosome regulation. Overexpression of HA-RAB12WT-K42A or HA-RAB12WT-Y111A was unable to induce centrosome expansion in astrocytes in contrast to the HA-RAB12WT as shown in the staining with anti-Pericentrin antibody (**Fig. 4e-f**). In addition, overexpression of HA-RAB12WT caused a smaller effect in the increasing volume of Pericentrin+ centrosomes in *Lrrk2* KO astrocytes than that in control astrocytes (**Fig. 4g-h, Fig. S6c**), suggesting that the effect of RAB12WT overexpression in centrosome amplification depends on LRRK2 and requires LRRK2 binding. Taken together, the above results demonstrated the requirement of a direct LRRK2 binding for the role of RAB12 in controlling the homeostasis of primary cilia and centrosomes.

#### RAB12-mediated recruitment of pRAB10/RILPL1 to centrosomes and elevation of pRAB10 levels requires LRRK2 binding in astrocytes

Previous studies suggested that RAB10 and RILPL1 regulate centrosomes and primary cilia^16, 18, 19, 24, 41^. We next asked whether RAB10 and RILPL1 participate in the action of RAB12 (or active RAB12) in the regulation of centrosome and cilia in astrocytes. Overexpression of HA-RAB12 promoted co-localization and clustering of the endogenous LRRK2 (**Fig. 5a)**, phospho-RAB10 (pRAB10) (**Fig. 5b**), and RILPL1 (**Fig. 5c**) with a marked increase of their staining intensity, compared to GFP-overexpressed astrocytes. The specificity of the endogenous LRRK2 staining was validated using *Lrrk2* KO astrocytes (**Fig. S7a**). The clustering of the endogenous LRRK2, pRAB10, and RILPL1 colocalized with Pericentrin or γ-tubulin staining (**Fig. 5a-c**), suggesting that they are recruited to amplified centrosomes or centrioles. Interestingly, although the overexpression of HA-RAB12QL also increased the intensity of pRAB10 and RAB12-LRRK2 co-clustering, the active mutant did not show a significant colocalization with pRAB10, RILPL1 or centrosome markers (**Fig. 5b-c**).

**Fig. 5.**
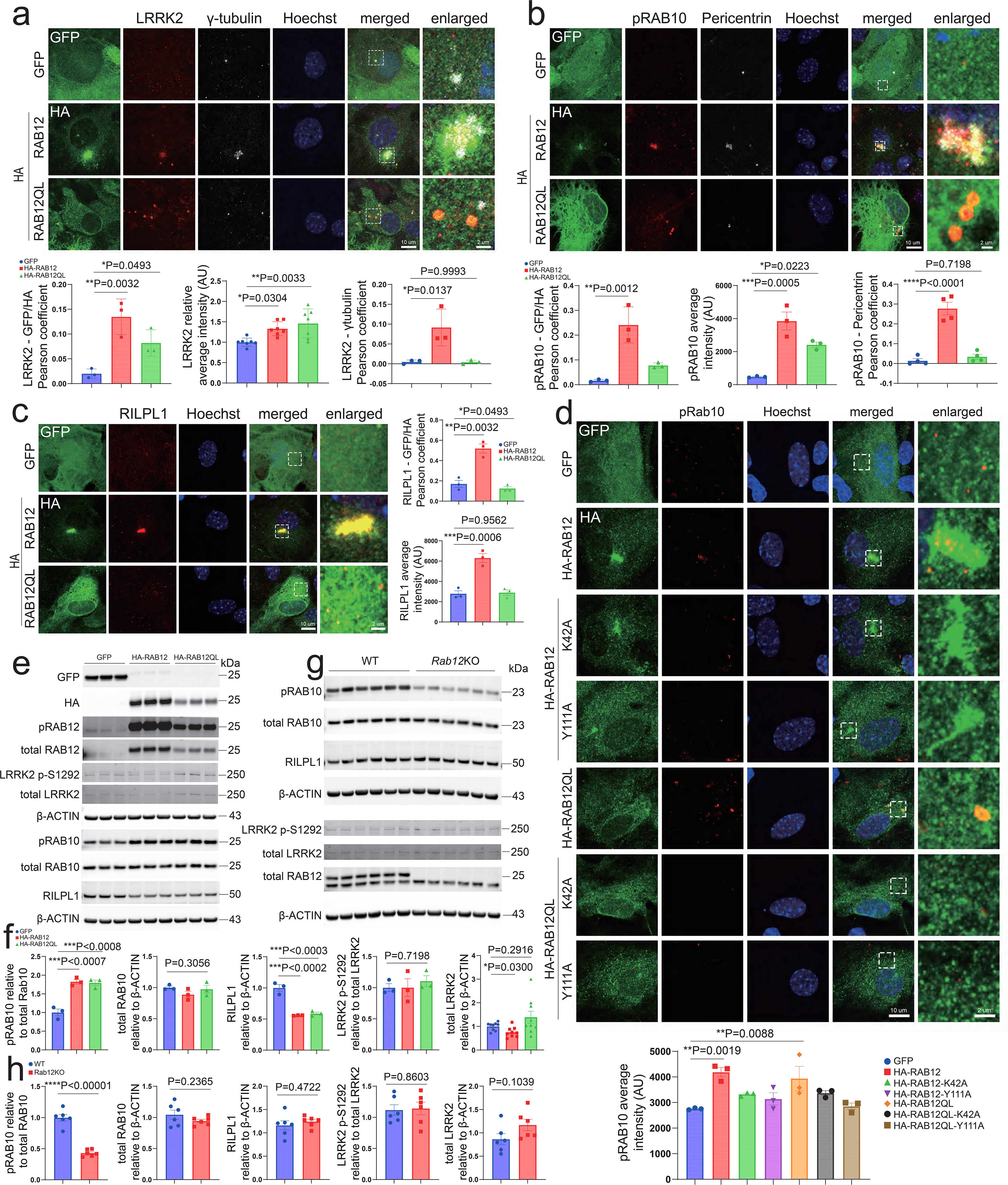
**Examination of LRRK2, pRAB10, and RILPL1 in astrocytes overexpressing RAB12. a**-**c**, Primary culture of astrocytes was infected with AAV-*GFAP*-*Gfp*, AAV-*GFAP*-HA-*Rab12*WT, or AAV- *GFAP*-HA-*Rab12*QL. IF staining was performed to detect LRRK2 (**a**), pRAB10 (**b**), and RILPL1 (**c**) in the cells by corresponding antibodies. Images as in (**a**-**c**) were quantified to determine LRRK2-GFP/HA Pearson coefficient, LRRK2 intensity colocalized with GFP or HA, and LRRK2-γ-tubulin Pearson coefficient in (**a**), pRAB10-GFP/HA Pearson coefficient, pRAB10 intensity colocalized with GFP or HA, and pRAB10-γ-tubulin Pearson coefficient in (**b**), and RILPL1-GFP/HA Pearson coefficient and RILPL1 intensity colocalized with GFP or HA in (**c**). **d**, AAVs carrying *Gfp*, HA-*RAB12*WT, HA-*RAB12*QL, or their LRRK2 binding mutants, such as HA-*RAB12*WT-K42A, HA-*RAB12*WT-Y111A, HA-*RAB12*QL- K42A, or HA-*RAB12*QL-Y111A were used to infect primary culture of astrocytes. IF staining was performed to detect pRab10 in the infected cells, and its intensity was measured. The magnified views of the dash boxes in the images in (**a**-**d**) were displayed in the last image column in each panel. **e**-**f**. Immunoblot analysis of primary cultured astrocytes overexpressing GFP, HA-RAB12WT, or HA- RAB12QL. The levels of pRAB10, total RAB10, RILPL1, LRRK2 p-S1292, total LRRK2, GFP, HA, pRAB12, and total RAB12 were detected. **g**-**h**, Immunoblot analysis of WT and *Rab12*KO primary cultured astrocytes. The levels of pRAB10, total RAB10, RILPL1, LRRK2 p-S1292, total LRRK2, pRAB12, and total RAB12 were quantified. Each dot in the graphs represents >60 cells from each mouse (**a-d**) and cells from an individual mouse (**f, h**). *P* values were calculated using one-way ANOVA followed by Turkey’s HSD post hoc tests (**a-d, f**) and unpaired two-tailed *t*-tests (**h**). \**P*<0.05, \*\**P*<0.01, \*\*\**P*<0.001, \*\*\*\**P*<0.0001; error bars = SEM; scale bars are indicated.

Furthermore, we found that overexpression of LRRK2-binding mutant HA-RAB12WT-K42A or HA- RAB12WT-Y111A failed to potentiate pRAB10 levels or recruit pRAB10 to the RAB12+ clusters (**Fig. 5d**). Lack of increase of pRAB10 intensity was also observed with the overexpression of active RAB12 mutants (HA-RAB12QL-K42A or HA-RAB12QL-Y111A) in astrocytes (**Fig. 5d**). In addition, the pRAB10 staining intensity was completely abolished in *Lrrk2* KO astrocytes with the overexpression of HA- RAB12WT or active HA-RAB12QL (**Fig. S7b-c**). The levels of RILPL1 colocalizing with HA-RAB12 was also markedly reduced in *Lrrk2* KO compared to the control astrocytes although the total fluorescent intensity of RILPL1 did not change (**Fig. S7d-f**), indicating RAB12 overexpression-induced RILPL1 recruitment is likely partially mediated by LRRK2.

We next performed immunoblot to analyze the impact of overexpression or loss-of-function of *Rab12* in astrocytes. Both WT and active RAB12 overexpression led to a significant increase of pRAB10 and reduction of RILPL1 and LRRK2 p-S935 levels, but with no impact on LRRK2 p-S1292 and total RAB10 levels. Only WT RAB12 overexpression, but not its active form, resulted in a reduction of endogenous total LRRK2 protein levels (**Fig. 5e-f, Fig.S7g-h**). In contrast, Rab12 KO astrocytes showed a remarkable decrease of pRAB10 and little change in the total RAB10, RILPL1, LRRK2 p-S935, p-S1292, or total LRRK2 levels (**Fig. 5g-h, Fig.S7i-j**). Thus, our results collectively demonstrate a critical role of *Rab12* in the control of RAB10 phosphorylation levels and the recruitment of pRAB10/RILP1 to centrosomes, which requires the direct LRRK2 binding.

#### Block of LRRK2 kinase activity prevents RAB12-mediated centrosome amplification and suppression of primary ciliogenesis

We next asked if the block of LRRK2 activity influences the RAB12-regulated centrosome and primary cilia homeostasis. We treated GFP-, HA-RAB12WT*-,* or HA-RAB12QL overexpressed astrocytes with a well-characterized LRRK2 inhibitor MLi2^42^ and found effective depletion of pRAB10 signals in all cases (**Fig. 6a-b**). The LRRK2 inhibitor significantly reduced the volume of γ-tubulin or Pericentrin labeled centrosomes in HA-RAB12WT overexpressed astrocytes (**Fig. 6c-e**), while it had a negligible effect on the centrosomes in GFP overexpressed astrocytes. Moreover, the MLi2 treatment rescued the decreased ciliated astrocyte percentage caused by the overexpression of active HA-RAB12QL and did not change the cilia length or volume (**Fig. 6f-i**). Thus, the data suggests that the block of LRRK2 activity abolishes the phosphorylation of RAB10 and consequently prevents the deregulated centrosome and primary ciliogenesis caused by RAB12 hyperactivity.

**Fig. 6.**
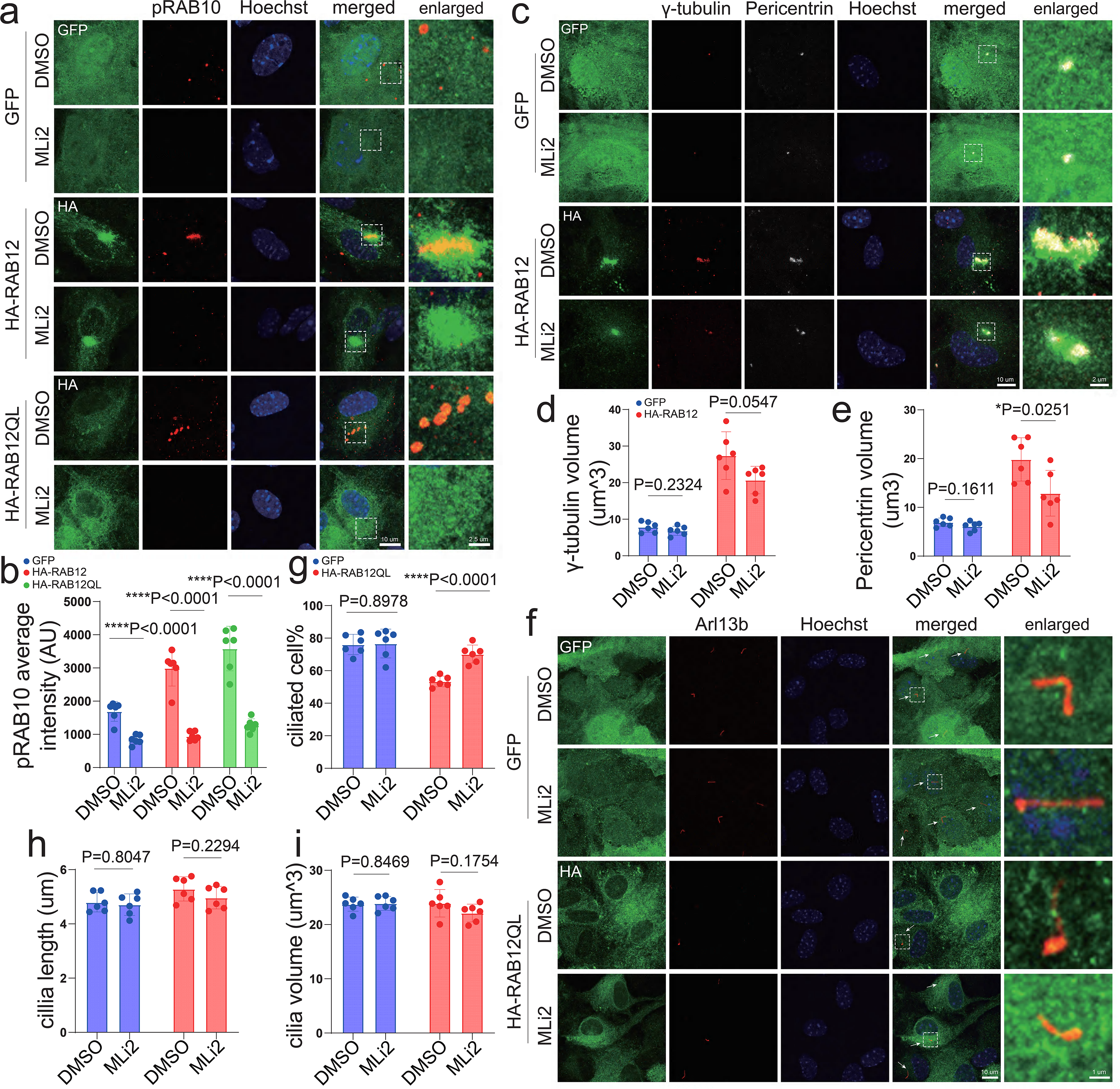
**The impact of LRRK2 kinase activity on RAB12’s cilia formation and centrosome homeostasis effects. a**-**b**, Primary culture of astrocytes was infected with AAV-*GFAP*-*Gfp*, AAV- *GFAP*-HA-*Rab12*WT, or AAV-*GFAP*-HA-*Rab12*QL and pRAB10 average intensity was quantified in the infected cells treated with MLi2 or DMSO. **c**-**e**, The volume of γ-tubulin and Pericentrin was examined in astrocytes infected with AAV-*GFAP*-*Gfp*, or AAV-*GFAP*-HA-*Rab12*WT and treated with MLi2 or DMSO. **f**-**i**, The ciliated cell percentage, cilia length, and cilia volume were quantified in astrocytes infected with AAV-*GFAP*-*Gfp*, or AAV-*GFAP*-HA-*Rab12*QL. The boxed areas in (**a**, **c**, **f**) were enlarged as shown. Each dot in the graphs represents >60 cells from each mouse. Unpaired two-tailed *t*-tests, \**P*<0.05, \*\*\*\**P*<0.0001; error bars = SEM; scale bars are indicated.

#### Loss of *Rab12* rescues *Lrrk2-*G2019S - induced ciliary deficiency and centrosome alteration in astrocytes

A previous study showed that PD mutation LRRK2-G2019S led to cilium deficiency and enhanced rate of split centrosomes due to its increased kinase activity^17^. We thus asked if the pathogenic effect of LRRK2-G2019S occurs in astrocytes and depends on RAB12. Indeed, we observed that the primary astrocytes from BAC transgenic *Lrrk2*-G2019S mice^32^ (GStg) showed decreased ciliated astrocyte percentage (with no change in cilia volume or length) and increased incidence of split centrosomes compared to control astrocytes (**Fig. 7**), consistent with the previous observations^24^. Interestingly, mutant astrocytes derived from compound mice combining GStg and *Rab12* KO showed no difference in the percentage of ciliation or split centrosomes when compared to control astrocytes (**Fig. 7**). The above observations demonstrated that the disruption of *Rab12* abolished the impact of the *Lrrk2* disease mutant on primary cilia and centrosome homeostasis in astrocytes.

**Fig. 7.**
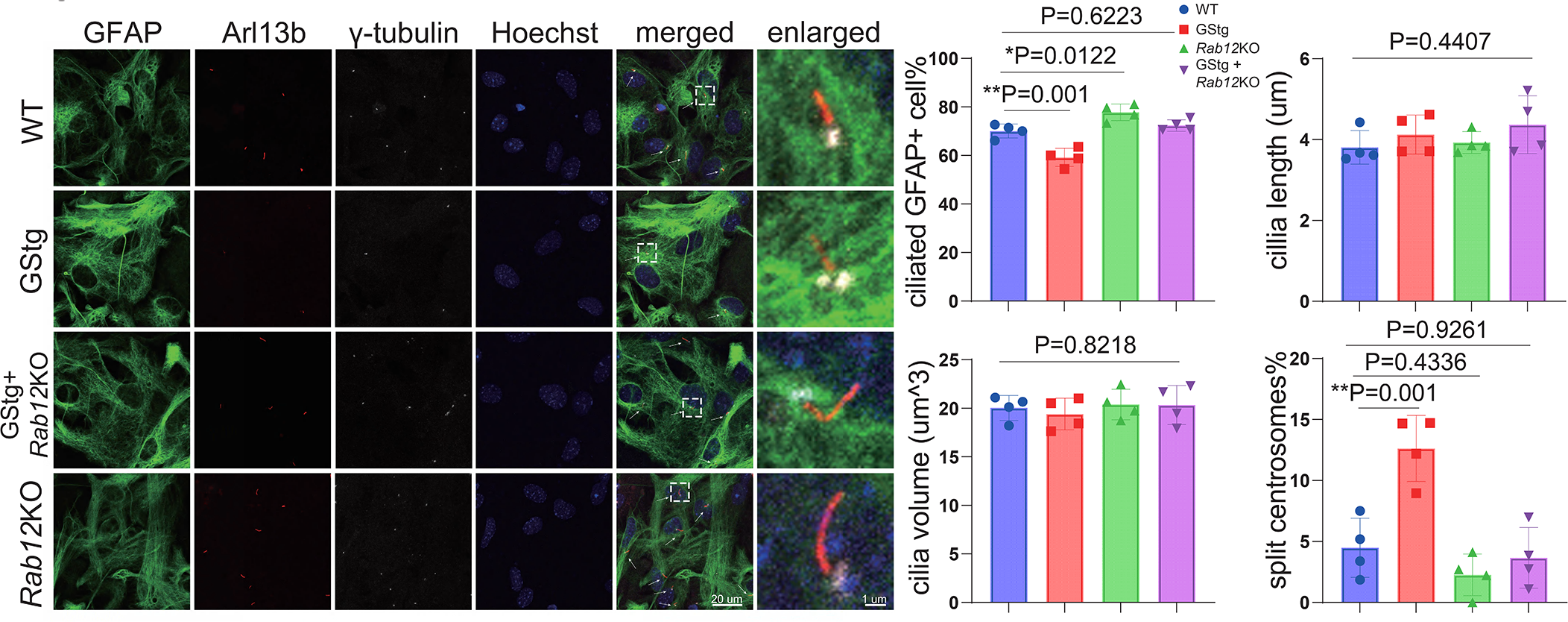
The effect of RAB12 on the ciliogenesis and centrosome alteration caused by *Lrrk2- G2019S* in astrocytes. The mice overexpressing *Lrrk2-*G2019S (GStg) were crossed with *Rab12* KO mice to generate GStg + *Rab12* KO double mutant mice. IF staining was performed to detect the cilia and the centrosomes in the primary cultures of astrocytes from WT, GStg, GStg + *Rab12* KO, and *Rab12* KO neonatal mice. The images from each group were analyzed to quantify the ciliated cell percentage, cilia length, cilia volume, and split centrosome percentage. In this study, split centrosomes are defined as the distance between the duplicated centrosomes in individual cells > 0.8 um^24, 64^. The enlarged images show the representative cilia and centrosomes inside the box areas. Each dot in the graphs represents >60 cells from each mouse. One-way ANOVA followed by Turkey’s HSD post hoc tests, \**P*<0.05, \*\**P*<0.01; error bars = SEM; scale bars are indicated.

## Discussion

Our current study determines the structure of RAB12–LRRK2 complex and provides the atomic details for RAB12–LRRK2 interaction. We show that the RAB12–LRRK2 complex regulates primary ciliogenesis and centrosome homeostasis in astrocytes and that RAB12 is crucial for LRRK2 kinase activity acting upon the phosphorylation of RAB10. Furthermore, pathogenic effects of LRRK2-G2019S on cilia and centrosome homeostasis in astrocyte depend on RAB12. Thus, our study identifies the interaction between LRRK2 and RAB12 as a new therapeutic target.

Our structural analysis demonstrates that multiple small RAB GTPases, including RAB12, RAB29, and RAB32a, share the similar binding site of the CDR1 and Switch I-Interswitch-Switch II motifs in LRRK2 binding ^34^. However, RAB12 differs from others by binding to the H1 helices of ARM6-ARM7, while RAB29 and RAB32 primarily interact with the H2 helices of ARM9-ARM10 of LRRK2 (**Fig. 1**). RAB12 and RAB29 were previously shown to activate LRRK2 when monitoring p-RAB10 and LRRK2 Ser1292 autophosphorylation^22, 31, 43^. The RAB29-dependent activation of LRRK2 with Ser1292 autophosphorylation was proposed to be mediated by the formation of asymmetric LRRK2 tetramers^34^. However, the binding of RAB12 on ARM6-ARM7 is not comparable with the tetrameric configuration of LRRK2 (**Fig. S3g**), suggesting that RAB12-dependent activation should be mechanistically different from RAB29-dependent activation of LRRK2 at the molecular level. In fact, we observed that deletion or overexpression of *Rab12* has little effect of Ser1292 autophosphorylation of LRRK2 in contrast to the phospho-Rab10 levels, indicating that the role of RAB12 in activating LRRK2 kinase activity could be substrate specific. It is worth noting though that the overexpression of active RAB12 may stabilize LRRK2 protein (**Fig. 5e**).

Through both loss-of-function and gain-of-function analysis, we establish the physiological function of RAB12 as an inhibitor of primary ciliogenesis in astrocytes. We show that RAB12 is essential for the maintenance of primary ciliary homeostasis in astrocytes of multiple brain regions including the cortex and the striatum. Our result indicates altered cilium size (unchanged ciliated cell percentage) in *Rab12* KO neurons, implicating RAB12 in ciliary regulation also in neurons. A recent study suggested a potential function of RAB12 in ciliogenesis in RPE and A549 cell cultures with unclear mechanism^22^. Our study provides an insight into the mechanism whereby RAB12 suppresses ciliogenesis. We observe that the expression of active RAB12 reduces ciliated astrocyte percentage, which requires a direct binding between RAB12 and LRRK2 and kinase activity of LRRK2. Interestingly, we find that overexpression of wildtype RAB12 not only abolishes the primary ciliogenesis, but also drives centrosome/centriole amplification. Such an effect of RAB12 also requires a direct LRRK2 interaction and LRRK2 kinase activity. Furthermore, the centrosome/centriole amplification involves the recruitment of p-RAB10 and RILPL1. The lack of centrosome phenotypes in astrocytes with active RAB12 overexpression may reflect an advanced stage, whereas wildtype RAB12 overexpression causes an intermediate phase that “arrest” amplified centrosomes to be resolved by RAB12 activation associated with membrane. Several lines of evidence have demonstrated that centrosome/centriole amplification could lead to the failure of ciliogenesis^37, 44, 45^. Thus, our study suggests a model that RAB12-LRRK2 protein complex suppresses primary ciliogenesis through altering centrosome/centriole homeostasis and recruiting centrosomal p- RAB10 and RILPL1. Previous works proposed that LRRK2 signaling converged on a centriolar p-RAB10- RILPL1 complex to cause deregulation of centrosomes and that ciliogenesis and centrosomal defects are the two distinct consequences of LRRK2 downstream signaling^24^. Our study, however, reveals RAB12 as a critical link coupling the two processes regulated by LRRK2 signaling. Furthermore, we observe no effect of RAB8a, RAB10, or LRRK2WT overexpression in altering primary cilia morphology or centrosome amplification, suggesting that RAB12 is a rate limiting factor.

Our study clearly demonstrates the physiological function of RAB12 in controlling homeostatic LRRK2 kinase activity towards the phosphorylation of small GTPase RAB10 in astrocytes or mouse brains, placing RAB12 upstream of LRRK2-RAB10 signaling. Our study also suggests the presence of RAB12- LRRK2 complex under basal condition, which helps maintain the homeostatic levels of phospho-RAB10. We hypothesize that the basal RAB12-LRRK2 complex plays a role in preventing excessive primary ciliogenesis through regulating centrosome homeostasis in astrocytes. Such a homeostatic function of RAB12-LRRK2 complex, however, is disrupted by the LRRK2-G2019S mutation. This gain-of-function mutation exacerbates the suppression of primary ciliogenesis and increases the occurrence of split centrosomes, based on our observations and others^19, 24^. It remains possible that the split centrosomes corroborate the stage of cell division in astrocytes, leading to the inhibition of primary ciliogenesis^24, 46^. Our study also demonstrates that the LRRK2-G2019S-induced defects in percentage of ciliation and split centrosomes were brought to the normal levels as in WT astrocyte after *Rab12* deletion (**Fig. 7**). The result reveals the critical role of RAB12 in mediating the pathogenic functions of the LRRK2-G2019S in impairing ciliogenesis and causing excessive split centrosomes in astrocytes. Our findings, therefore, suggest that disruption of RAB12-LRRK2 interaction can be explored to block LRRK2 pathogenic signaling in addition to LRRK2 kinase inhibition in therapeutic development. For such a purpose, our RAB12-LRRK2 structure provides valuable information for designing the intervention to disrupt the RAB12-LRRK2 complex formation.

Our study has limitations and raises many questions. First, we showed that RAB12 is a robust substrate of LRRK2 in the brain and forms a complex with LRRK2. It is unclear, however, how phosphorylation of RAB12 may affect RAB12-LRRK2 interaction and RAB12 function. According to the structure of the RAB12-LRRK2 complex, RAB12’s phosphorylation site S106 does not directly involve binding to LRRK2 (**Fig.1b**). We speculate that the p-S106 of Rab12 is unlikely to affect its binding to LRRK2. A report also showed that RAB12’s phospho-dead mutant is fully capable of recruiting and activating LRRK2 ^22^. Future studies should elucidate the function of RAB12 phosphorylation. Second, our phospho-proteomics study did not find any significant changes in the phosphorylation sites of known LRRK2 substrates (RAB proteins) beyond RAB12, such as RAB8, RAB10, or RAB29. It remains unclear whether RAB12 is the most (patho)physiological substrate of LRRK2 in the brains. The result could be due to the low abundance of the above RAB proteins in mouse brain tissues and sensitivity of the current methodology. Interestingly, a previous report found that LRRK2-R1441C mutation increases the levels of pRAB12, but not pRAB10, in mouse brain, lung, and kidney tissues^47^. Third, the detailed molecular mechanism for how RAB12- LRRK2 complex inhibits ciliogenesis and regulates centrosome homeostasis is unknown.

Finally, although our study establishes that RAB12-LRRK2 complex controls ciliary and centrosomal homeostasis, it is unclear whether such a function is related to the previously reported role of RAB12 in regulating lysosome damage or stress response^29, 31^. Furthermore, the significance of astrocytic ciliary defects and centrosome abnormality induced by the LRRK2 variant in the pathogenesis of PD remains to be clarified. Nonetheless, our structural analysis and functional characterization of RAB12-LRRK2 complex provides an insight into the pathogenic mechanism and paves the way for the development of novel therapeutics for PD.

## Methods

### 1. Cloning, expression and purification of human LRRK2 and RAB12

Human LRRK2 construct was expressed and purified as described^4, 34^. Briefly, bacmids carrying LRRK2 construct was generated in E. coli DH10Bac cells (Invitrogen). Recombinant baculoviruses were produced and amplified in Sf9 cells. Then P3 virus of full-length LRRK2 was used to transfect HEK293F cells for protein expression. Briefly, for 1 L cultures of HEK293F cells (∼2-3x106 cells/mL) in Freestyle 293 media (Gibco) supplemented with 2% FBS (Gibco), about 100 ml P3 virus was used.

Infected cells were incubated at 37℃ overnight, and protein expression was induced by adding 10 mM sodium butyrate. Cells were cultured at 30℃ for another 48-60 h before harvest. The cell pellet from the 600 mL culture was resuspended in 30 mL lysis buffer (20 mM Tris pH 8.0, 200 mM NaCl, 5% glycerol, 2 mM DTT and protease inhibitors), and then cells were lysed by brief sonication. LRRK2 was separated from the insoluble fraction by high-speed centrifugation (38,000 × g for 1 h), and incubated with 1 mL CNBr-activated sepharose beads (GE Healthcare) coupled with 1 mg high-affinity GFP nanobodies (GFP-NB)^48^. The GFP tag was cleaved by preScission protease at 4℃, and LRRK2 was further purified by size-exclusion chromatography with a Superose 6 Increase 10/300 GL column (GE Healthcare) equilibrated with 20 mM Tris pH 8.0, 200 mM NaCl and 2 mM DTT. The purified protein was collected and concentrated to 12.5 mg/ml (OD280) using a 100-kDa MWCO centrifugal device (Ambion), flash-frozen in liquid N2 and stored at -80℃.

DNA encoding human RAB12 (residues 36-211) was synthesized by Genewiz and subcloned into the modified pGEX plasmid (Sigma), which attaches a GST tag and a TEV protease cleavage site at the N-terminus. For structural and biochemical studies, we generated RAB12Q101L mutant, which was constructed using the QuickChange Site-Directed Mutagenesis kit (Strategene). The recombinant proteins were overexpressed in E. coli strain BL21 (DE3) in LB media supplemented with 0.05 mg/ml ampicillin, and cells were grown at 37℃ until OD600 reached 0.8. Further expression was induced by adding 0.4 mM IPTG, and cells were allowed to grow at 16℃ for 20 h. Harvested cells were lysed by sonication in the lysis buffer (20 mM Tris-HCl pH 8.0, 200 mM NaCl, 5% glycerol, 5 mM MgCl2, and 1 mM PMSF), and lysates were cleared by centrifugation at 38,000 × g for 45 min. Subsequently, the proteins were purified by GST affinity chromatography on the glutathione Sepharose beads (GE Healthcare) in lysis buffer. The bound proteins were then eluted with lysis buffer containing 20 mM GSH and incubated with 1 mM GppNHp at 4℃ overnight, followed by gel filtration chromatography using a Superdex 200 Increase 10/300 GL column (GE Healthcare) in a storage buffer (20 mM Tris-HCl pH 8.0, 100 mM NaCl, 1 mM MgCl2, and 2 mM DTT). For the structural study, the GST tag was removed by on-column cleavage with TEV protease at 4℃ overnight before gel filtration chromatography. The peak fractions were collected and concentrated to 4.3 mg/ml (OD280), flash-frozen in liquid N2, and stored at -80℃.

### 2. Cryo-EM sample preparation

Cryo-EM grids were prepared with a Vitrobot Mark IV (FEI). Quantifoil R1.2/1.3 300 Au holey carbon grids (Quantifoil) were glow-discharged for 30 s. Purified full-length LRRK2 and RAB12Q101L were incubated together on the ice for 1 h with a final concentration of 30 mM and 60 mM in the presence of 2 mM AMP-PNP, 1 mM GppNHp and 1 mM MgCl2. In addition, 2.3 mM fluorinated Fos-Choline-8 was added right before freezing the grids, and then 3.5 mL of protein sample were pipetted onto the grids, which were blotted for 3.5 s under blot force -3 at 16℃ and 95% relative humidity and plunge-frozen in liquid nitrogen-cooled liquid ethane.

### 3. Cryo-EM data acquisition and processing

The RAB12–LRRK2 dataset was collected on a Titan Krios (Thermo Fisher Scientific) transmission electron microscope equipped with a K3 direct electron detector and post-column GIF energy filter (Gatan). Data collection was performed in an automated manner using EPU (Thermo Fisher Scientific). Movies were recorded at defocus values from -1.2 to -2.6 mm at a magnification of 130kx in hardware binning mode, corresponding to a pixel size of 0.6485 Å at the specimen. During 2.0 s exposure, 70 frames were collected with a total electron dose of ∼78 e-/Å-2 (at a dose rate of 1.1143 e-/frame/Å2). In total, 20,023 images were collected. Motion correction was performed on hardware-binned movie stacks and binned by 1 using MotionCor2^49^. Contrast transfer function (CTF) estimation was performed using Gctf^50^. Prior to particle picking, micrographs were analyzed for good power spectrum, and the bad ones were discarded (with 19,637 good images remaining).

Particles were selected using the template picker with reported inactive full-length LRRK2 structures^4^ as a reference in cryoSPARC^51^ and extracted using a binning factor of 2. Several rounds of the 2D classification were performed to eliminate ice artifacts, carbon edges, and false-positive particles containing noise. During 2D classification, two groups of good classes were observed, corresponding to the RAB12–LRRK2 monomer and dimer states, respectively. Both groups were selected, and ab initio reconstruction was performed. In order to further separate the RAB12–LRRK2 monomer and dimer states, we performed Heterogeneous refinement in cryoSPARC. As a result, 95,596 particles were assigned to the monomer class and 71,784 particles to the dimer. Both 3D classes were further refined using cryoSPARC after extraction of unbinned particles corresponding to each identified sub-set. For the RAB12–LRRK2 monomer state, we performed a standard NU-refinement without imposing symmetry, followed by a focused 3D refinement (with a soft mask around RAB12–LRRK2 interface region) to improve the map quality of the ARM domain of the LRRK2 with RAB12 binding. All resolution estimates were calculated according to the gold-standard Fourier shell correlation (FSC) using the 0.143 criterion^52^. Local resolution was estimated in cryoSPARC. The density maps were B-factor sharpened in cryoSPARC and used to produce figures and build model.

### 4. Model building and refinement

The reported structures of LRRK2 (PDB 7LI4)^4^ and RAB12 (PDB 2IL1) were fitted and adjusted into the cryo-EM maps of RAB12–LRRK2 monomer state using Chimera^53^ and Coot^54^. The structural model was refined against the map using the real space refinement module with secondary structure and non-crystallographic symmetry restraints in the Phenix package^55^. Fourier shell correlation curves were calculated between the refined model and the full map. The geometries of the models were validated using MolProbity^56^. All the figures were prepared in PyMOL (Schrödinger, LLC.), UCSF Chimera^53^ and UCSF ChimeraX^57^.

### 5. Animals

All animal studies were carried out according to the guidance of the Icahn School of Medicine at Mount Sinai Animal Care and Use Committee (IACUC-2021-00031). All the mice in this study were adopted under a 12 h light/12 h dark cycle environment at 20-23 ℃ and 40-60% humidity with available food and water. *Lrrk2* BAC transgenic mice overexpressing *Lrrk2* wildtype (WTtg) and PD-mutant G2019S (GStg) were generated by our group^32^. *Lrrk2* knockout (KO) mice (Strain #: 016121) and C57BL/6J mice (WT, Strain #: 000664) were purchased from The Jackson Laboratory. *Rab12* KO mice were generated by deleting the exon 3 of *Rab12* gene from the mouse genome of C57BL/6J background using CRISPR-Cas9 technology by our group. For the generation of GStg + *Rab12* KO double mutant mice, GStg heterozygous mice were crossed with *Rab12* KO homozygous mice to obtain GStg heterozygous + *Rab12* KO homozygous mice. PCR was performed with genomic DNA isolated from mouse tail clips to confirm mice genotypes.

### 6. Mouse brain tissue collection and processing

Mice were anesthetized with 10 mg/kg xylazine and 50 mg/kg ketamine by intraperitoneal injection. For the samples used for phosphoproteome profiling and western blot, the mice were sacrificed by cervical dislocation after anesthetization. Then, striatal tissues were dissected from their brains and stored at – 80 ℃ for further processing. For the samples used for immunofluorescent staining, the anesthetized mice were cardiac perfused with cold PBS followed by 4% paraformaldehyde-PBS solution. The mouse brains were dissected, kept in the 4% paraformaldehyde-PBS solution at 4℃ overnight, and sequentially dehydrated in 30% sucrose-PBS buffer for 72 h. Finally, the brain tissues were embedded in OTC (Fisher Scientific, #23-730-571), frozen on dry ice, and sectioned at 30 ums with a cryostat (Leica) for the following experiments.

### 7. Whole proteome profiling and phosphoproteome profiling by TMT-LC/LC-MS/MS

Proteome and phosphoproteome profiling were performed as described previously^58^. Mouse brain samples were lysed in fresh lysis buffer (50 mM HEPES, pH 8.5, 8 M urea, 0.5% sodium deoxycholate and phosphatase inhibitor cocktail (PhosphoSTOP, Roche). The protein concentration of the lysate was quantified by BCA protein assay (Thermo Fisher Scientific). One mg of proteins from each sample were digested with Lys-C (Wako, 1:100 w/w) at room temperature for 2 h, diluted 4 times with 50 mM HEPES, pH 8.5, and further digested with trypsin (Promega, 1:50 w/w) overnight at room temperature followed by Cys reduction and alkylation. The peptides were acidified by 1% trifluoroacetic acid, desalted with Sep-Pak C18 cartridge (Waters), and then labeled with 10-plex TMT reagents (Thermo Fisher Scientific) followed by mixing equally. The mixture was desalted, dried and solubilized in 600 µL of buffer A (10 mM ammonium formate, pH 8) and separated on two concatenated XBridge C18 columns (3.5 μm particle size, 4.6 mm x 25 cm, Waters) into 42 fractions with a 60 min gradient from 13% to 45% buffer B (95% acetonitrile, 10 mM ammonium formate, pH 8) with flow rate of 0.4 mL/min. For each fraction, phosphopeptide enrichments were carried out by TiO_2_ beads (GL sciences) as previously reported^59^. Briefly, the dried peptides were dissolved in 100 μL of binding buffer (65% acetonitrile, 2% TFA, and 1 mM KH2PO4). TiO2 beads (1 mg) were washed twice with washing buffer (65% acetonitrile, 0.1% TFA), mixed with the peptide solution, and incubated with end-over-end rotation at room temperature for 20 min. The phosphopeptide-bound beads were collected by brief centrifugation, washed twice with 150 μL washing buffer and transferred to a C18 StageTip (Thermo Fisher Scientific) sitting on the top of a 2-mL centrifuge tube. The StageTip was centrifuged to remove the wash buffer completely and phosphopeptides were eluted under basic pH condition (15 μL, 15% NH4OH, 40% acetonitrile). The eluents were dried, dissolved in 5% formic acid and loaded on a reverse phase column (50 µm x 40 cm, 1.9 µm C18 resin (Dr. Maisch GmbH, Germany) interfaced with a Q-Exactive HF mass spectrometer (Thermo Fisher Scientific). Peptides were eluted at 65 °C by 9- 35% buffer B gradient in 4 h (buffer A: 0.2% formic acid, 3% DMSO; buffer B: buffer A plus 67% acetonitrile, flow rate of 0.15 µL/min). The mass spectrometer was operated in data-dependent mode with a survey scan in Orbitrap (60,000 resolution, 1×10^6^ AGC target and 50 ms maximal ion time) and 20 MS/MS high resolution scans (60,000 resolution, 1x10^5^ AGC target, 105 ms maximal ion time, HCD, 35 normalized collision energy, 1.5 m/z isolation window, and 20 s dynamic exclusion). The whole proteome profiling was performed in a similar protocol but excluding the step of phosphopeptide enrichment.

The data analysis was performed by our JUMP suite^60^ Briefly, acquired MS/MS raw files were converted into mzXML format and searched by the JUMP algorithm against a composite target/decoy database to estimate FDR. The target protein database was downloaded from the Uniprot mouse database and the decoy protein database was generated by reversing all target protein sequences.

Searches were performed with 10 ppm mass tolerance for precursor ions and 15 ppm for fragment ions, fully tryptic restriction, two maximal missed cleavages and the assignment of a, b, and y ions. TMT tags on lysine residues and peptide N termini (+229.16293 Da) and carbamidomethylation of Cys residues (+57.02146 Da) were used for static modifications and the dynamic modifications included the oxidation of methionine residues (+15.99492 Da) and Ser/Thr/Tyr phosphorylation (+79.96633). The assigned peptides were filtered by mass accuracy, minimal peptide length, matching scores, charge state and trypticity to reduce phosphopeptide FDR to 1%. Phosphosite reliability was evaluated by the localization score (Lscore) from JUMP suite. TMT reporter ion intensities of each PSM were extracted. The raw intensities were then corrected based on isotopic distribution of each labeling reagent and loading bias. The mean-centered intensities across samples were calculated and phosphopeptide intensities were derived by averaging related PSMs. Finally, phosphopeptide absolute intensities were determined by multiplying the relative intensities by the grand-mean of three most highly abundant PSMs. The whole proteome analysis was carried out similarly but excluding the mass shift of phosphorylation.

Proteins or phosphopeptides showing changes were identified using *P* values calculated through the limma package^61^, which employs moderated *one-way ANOVA* for the four genotypes’ comparison.

These *P* values were subsequently adjusted using the *Benjamini-Hochberg* procedure to control the false discovery rate. Phosphopeptides that passed the adjusted *P* value threshold of 0.05 were then examined manually. In addition, the phosphopeptide levels were also normalized by their related protein levels to investigate the impact of kinase activity^62^.

### 8. Plasmid construction

Total RNA was purified from the mouse brain tissue using RNeasy Mini Kit (QIAN, #74106) and was reverse-transcribed to cDNA. The mouse *Rab12*WT (m*Rab12*WT) coding sequence was amplified from the cDNA obtained. An HA tag was added to the N terminal of the m*Rab*12WT coding sequence and subcloned into pAAV-*GFAP-EGfp* (addgene, #50473) using SalI and EcoRI restriction enzyme sites and LV-*hGFAP-Gfp* (addgene, #183906) using AgeI and XhoI restriction enzyme sites, respectively.

The original DNA fragments between the restriction enzyme sites in the two plasmids were removed. Mouse *Rilpl1*, *Rab8a*, and *Rab10* were also amplified from the cDNA above. *Rilpl1* was conjugated with a His tag at its C terminal, and a V5 tag and a Flag tag were conjugated at the N terminal of *Rab8a* and *Rab10*, respectively. All three genes were also inserted into LV-*hGFAP-Gfp* using AgeI and XhoI sites. Human *RAB12*WT (h*RAB12*WT) and its QL variant (h*RAB12*QL) DNA fragments were amplified from pcDNA3.1-h*RAB12*WT and pcDNA3.1-h*RAB12*QL generated in our lab before. NEBuilder® HiFi DNA Assembly Kit (NEB, # E2621L) was used to construct the m*Rab12*Q100L variant, h*RAB12*WT and h*RAB12*Q101L binding mutations (K42A, F79A, R92A, and Y111A). All these mutant variants were subcloned into the same location of pAAV-*hGFAP-GFP* as done with mRAB12WT.

### 9. Virus production

For AAV particles generation for *in-vitro* studies, the AAV plasmids harboring the genes of interest were co-transfected with helper plasmids pAAV2/1 (addgene, #112862) and pAdDeltaF6 (addgene, #112867) into HEK293T cells using lipofectamine 3000 (Thermo Fisher Scientific, #L3000008). The medium was changed 6 h after transfection and replaced with fresh culture medium (DMEM with 10% fetal bovine serum). The medium was collected 48 h after transfection and centrifuged at 1000 x g for 10 min to spin down the cell debris. The supernatant was transferred to a new tube followed by addition of Polyethylene glycol 8000 (PEG000)-NaCl buffer (40% PEG8000, 2.5M NaCl) to precipitate the AAV particles at 4 ℃ overnight. Then, the AAV particles were spun down at 4000 x g at 4 ℃ for 1 h. The supernatant was removed and the remaining pellets were resuspended in glia medium (DMEM with 10% foetal bovine serum, 1x GlutaMax, 1x sodium pyruvate, and 1% penicillin/streptomycin). The resuspended AAV particles were aliquoted and frozen at -80 ℃.

For lentivirus production, HEK293T cells were co-transfected with lentiviral expression plasmids containing the gene of interest and helper plasmids pMDL, pRSV, and pVSV-G. The culture was replaced with fresh culture medium 6 h post-transfection. The culture medium containing the lentivirus particles was collected at 24 h, 48 h, and 60 h post-transfection. The medium from the indicated time points was combined and centrifuged at 1000 x g for 10 min to remove the cell debris, and then transferred to a sterile ultracentrifuge tube. The lentivirus particles were precipitated by ultracentrifugation at 2,1000 RPM at 4 ℃ for 2 h. The supernatant was removed, and the lentivirus pellets were re-suspended with 1 x lenti-freezing medium (0.5 M sucrose in DMEM), aliquoted, and frozen at -80 ℃.

### 10. Primary astrocyte culture

Briefly, P0-P2 neonatal mice were anesthetized on ice. The whole brains were dissected, and the olfactory bulbs, meninges, cerebellums, and brain stems were removed. The remaining tissues were kept in tubes containing chilled glia medium composed of DMEM with 10% foetal bovine serum (HyClone, #SH30396.03HI), 1x GlutaMax (GIBCO, #35050-061), 1x sodium pyruvate (GIBCO, #11360- 070), and 1% penicillin/streptomycin (GIBCO, #15140-122). Brain tissue from different neonatal mice were placed in separate tubes. Cells were then dissociated using 1 mL pipette tips followed by a fire-polished glass pipette and cultured in T75 flasks in a humidified 5% CO2 incubator at 37°C for 14 days. On DIV14, the astrocytes were detached from the flasks’ bottoms by 0.05% trypsin (GIBCO, #25200056) and replaced on culture plates for either immunoblot analysis or immunofluorescent staining.

### 11. Immunofluorescent staining

For immunofluorescent staining of the mouse brain, brain slices were mounted on a superfrost plus microscope slide (Thermo Fisher Scientific, # 12-550-15) and washed with PBS once for 5 min. Slices were blocked with blocking buffer (5% goat serum in PBS containing 0.3% Triton X-100) for 1 h at room temperature and incubated with primary antibodies at 4 ℃ for about 16 h. Then, slices were rinsed with PBST (PBS containing 0.05% Triton X-100) twice for 5 min each and once with PBS for 5 min, followed by incubation with corresponding fluorescent dye-conjugated secondary antibodies for 1 h at room temperature shielded from light. The secondary antibody was rinsed from the slices as previously indicated and incubated with 2 uM Hoechst solution (Thermo Fisher Scientific, #62249) for 5 min at room temperature. Finally, slices were rinsed with PBS twice for 5 min and mounted with ProLong™ Diamond Antifade Mountant (Thermo Fisher Scientific, # P36961). For the immunofluorescent staining for primary cultured astrocytes, the cells were fixed with 4% paraformaldehyde-PBS solution for 15 min at room temperature, and sequentially washed three times with PBST. The cells were blocked and stained in a similar manner as mentioned above except the concentration of Triton X-100 in the blocking buffer was reduced to 0.1%. For the centrosome staining, the cells were additionally fixed with methanol at -20℃ for 15 min after paraformaldehyde fixation for enhanced visualization. The information on the antibodies used are listed in Table S5.

### 12. Co-immunoprecipitation

HEK293T cell line was maintained in DMEM supplemented with penicillin/streptomycin and 10% FBS (R&D Systems, #S11550). One day prior to transfection, HEK293T cells were seeded onto a 6-well plate at 6x10^5^ cells per well. Cells were transfected using Lipofectamine® 3000 (Invitrogen, #L3000015). Plasmid DNA concentrations of 0.5 ug were used for *RAB12*WT, *RAB12*WT-Y111A, *RAB12*WT-K42A, *RAB12*QL, *RAB12*QL-Y111A, *RAB12*QL-K42A plasmids and co-transfected with 4.5 ug of Flag tagged LRRK2WT plasmids for double transfected conditions. Plasmid DNA was added to 125uL Opti-MEM containing 10uL of P3000 followed by addition of 125 uL of Opti-MEM containing 3.75 uL Lipofectamine 3000. The plate was incubated at 37°C for 24 h in a 5% CO2 incubator. After incubation, transfected cells were then used for co-immunoprecipitation (Co-IP). Cells were lysed on ice with lysis buffer consisting of 1% Triton X-100 TBS buffer containing 1 x Halt™ protease and phosphatase inhibitor cocktail (#78440, Thermofisher Scientific) and 5 mM EDTA (Thermo Fisher Scientific #1861274). The concentration of cell lysates was examined by BCA assay (Thermo Fisher Scientificfi#23227) and 1000 ug of proteins from each sample were incubated in 30 uL Anti-FLAG® M2 Magnetic Beads (Millipore Sigma, #M8823) overnight at 4 ℃. After incubation, beads were washed three times with 1% Triton X-100 TBS. Proteins bound to the M2 Magnetic Beads were eluted by boiling in 1x loading buffers composed of NuPAGE™ LDS Sample Buffer (Thermo Fisher Scientificfi #NP0007) and 0.05M DTT (Sigmafi#11583786001) at 95 ℃ for 10 min. The supernatants were transferred into new Eppendorf tubes for immunoblot analysis analysis after brief centrifugation.

### 13. Immunoblot analysis

Brain tissue samples were homogenized through bead disruption using the Bullet Blender (Next Advance) technology. Samples were put in 1.5mL RINO tubes (Next Advance Inc.) with zirconium oxide beads (ZrOB05 + ZrOB10 mixture, Next Advance Inc) in the lysis buffer described in the Co-IP procedure but without Triton X-100 and homogenized for 2 min at speed 12 at 4 ℃. The tissue homogenates were mixed with equal volume of lysis buffer with 2% Triton X-100, and then lysed on ice for 20 min before centrifugation. For cell culture samples, cells were harvested and lysed in the lysis buffer on ice for 15 min, vortexed briefly before centrifugation. The centrifugation conditions for tissue samples and cell samples were both at 16000 x g for 10 min at 4 ℃. The supernatants were transferred to new tubes and BCA assay was performed to determine the samples’ protein concentrations. The samples were mixed with 3x loading buffers to a final concentration of 1x, and incubated at 70 ℃ for 10 min. After being cooled down on ice, the samples were loaded onto an SDS-PAGE gel. Proteins were separated by electrophoresis at 120 Volts for 1.5 h and then electroeluted onto a PVDF membrane (BIO-RAD, #10026933) at 25 Volts, 1 Ampere for 0.5 h using Trans-Blot® Turbo™ Transfer System (BIO-RAD). PVDF membranes were then washed with TBS for 5 min and blocked with Intercept Blocking Buffers (LI-COR, #927-60001) for 1 h at room temperature. Membranes were incubated in primary antibodies overnight at 4 ℃, followed by three times wash in TBST (0.1% Tween-20 in TBS) for 5 min each. The appropriate secondary fluorescent antibodies were applied for 1 h at room temperature and then washed off three times for 5 min each before the signals were visualized by the ChemiDoc Imaging system (BIO-RAD). Signal intensities were then measured by ImageJ (https://imagej.nih.gov/ij/index.html).

### 14. Electron microscope

Primary astrocytes were grown on Permanox® Slide (Electron Microscopy Sciences, #70400). Samples were prepared by washing cells with 0.1 M sodium cacodylate solution (Electron Microscopy Sciences, #15960-01) three times and fixed with sodium cacodylate solution containing 2% paraformaldehyde and 2.5% glutaraldehyde for 2 h at 4℃. They were then post fixed with sodium cacodylate containing 2% osmium tetroxide and 1.5% potassium ferricyanide for 1 h at room temperature. After being briefly washed by sodium cacodylate buffer, the cells were stained with 2% uranyl acetate in distilled H_2_O for 1 h at room temperature followed by dehydration in an ascending ethanol series. The cells were then embedded in resin through an increasing ethanol/resin series for 16-18 h (Electron Microscopy Sciences Embed 812 Kit, #14120), and incubated in a vacuum oven for 72 h at 60 ℃. Ultra-thin sections were obtained on copper 300 mesh grids (Electron Microscopy Sciences, G300H-Cu) and counterstained with 1% uranyl acetate and lead citrate. Images were taken by an HT7500 transmission electron microscope (Hitachi High-Technologies, Japan). Sample processing and imaging was performed by the Microscopy and Advanced Bioimaging CoRE at the Icahn School of Medicine at Mount Sinai.

### 15. Image acquisition and analysis

The fluorescent image acquisition in this study was performed using z-stack with an 63x objective at 1024 x 1024 resolution with the Zeiss LSM 780 confocal microscope (Carl Zeiss). The z-step interval was 0.5 um for astrocytes, and 0.75 um for neurons^63^. Sixteen z-stack images were taken for cultured astrocytes while 50 images for mouse brain slices. The images from astrocytes and neurons were obtained from the dorsal striatum and frontal cortex areas in the mouse brain slices.

For ciliated cell^15, 63^ and split centrosome^24, 64^ percentage quantitation, the z-stack images were collapsed by the maximum intensity projection function in ImageJ (https://imagej.nih.gov/ij/index.html) and manually counted as described previously. CiliaQ, an open-source ImageJ plugin^65^, was used to measure the cilia length and volume of primary cultured astrocytes, whereas the surface module in Imaris software (Bitplane) was utilized to determine the cilia length and volume of astrocytes and neurons in the mouse brain slices, and the centrosome volume of primary cultured astrocytes. The fluorescent intensity and Pearson coefficient between proteins were quantified by the particle analysis plugin and the JaCoP plugin^66^ in ImageJ, respectively.

### 16. Statistical analysis

GraphPad Prism 9 was used to execute the statistical analyses in this study. All the results were presented as mean ± standard error (SEM). As for comparisons between two groups, *Student’s t-test* was applied to determine the significance. As for the comparisons with more than two groups, *one-way ANOVA* was performed. *Turkey’s HSD post hoc test* was used for multiple group comparisons when *one-way ANOVA* indicated statistical significance. *P* value < 0.05 was defined as statistical significance.

## Acknowledgement

We thank the support of Electron Microscopy core facility at Mount Sinai and the Cryo-Electron Microscopy Center at St. Jude Children’s Research Hospital for help with cryo-EM data collection. We thank members of the Yue lab and members of the Sun lab for critical reading and helpful discussion and P. Hixson for cell-culture support. This work was funded by NIH (P20NS123320), The Michael J. Fox Foundation, and Parkinson’s Foundation to ZY. This work was also funded by the American Lebanese Syrian Associated Charities (ALSAC) and NIH (R01NS129795) to JS.

## Author contribution

Z.Y. conceived, designed, supervised the study and wrote the manuscript; J.S. conceived and supervised the structure research and wrote the manuscript; X-J.L. performed the cellular, molecular, and mouse experiments, analyzed the experimental data, and wrote the manuscript; H.W. conducted the structure experiments and related data analysis, and wrote the manuscript; B.T.H performed the Co-IP experiments, data analysis and wrote the manuscript; H.K. sectioned the mouse brains for the IF experiments and analyzed the experimental data; X-T.L. participated in establishing the mouse models; H.T. and J.P. carried out the phosphoproteomics and bioinformatics analysis and wrote the manuscript; I.C. carried out some of the culture experiments and analyzed the experimental data; Y.Z. performed some of the mouse genotyping experiments and analyzed the data. P.X. contributed financial support to the project and participated in the discussion in the manuscript preparation.

## Data and code availability

The cryo-EM maps of the RAB12–LRRK2 complex have been deposited in the Electron Microscopy Data Bank under the accession codes EMD-43234 (RAB12–LRRK2 monomer) and EMD-43235 (RAB12–LRRK2 dimer). The corresponding coordinates have been deposited in the Protein Data Bank under the accession codes 8VH4 (RAB12–LRRK2 monomer) and 8VH5 (RAB12–LRRK2 dimer). The mass spectrometry proteomics data have been deposited to the ProteomeXchange Consortium via the PRIDE partner repository with the dataset identifier PXD048732.

---

**Fig. S1.**
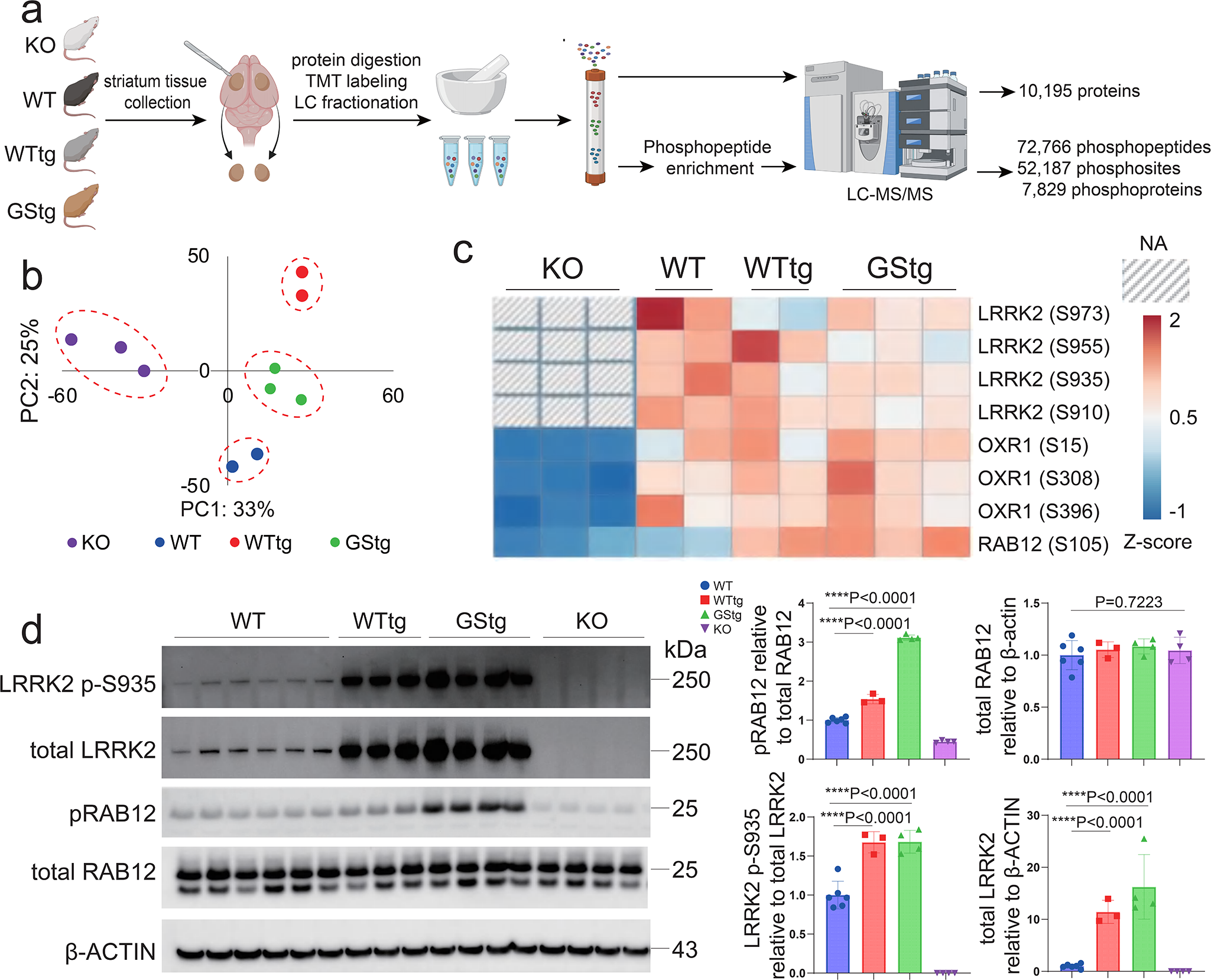
**Phosphoproteome analysis of LRRK2 kinase substrates in mouse brains. a**, Proteome and phosphoproteome profilings of mouse striata with *Lrrk2* KO (KO), WT, *Lrrk2* WT overexpression (WTtg), or *Lrrk2*-G2019S overexpression (GStg) were analyzed by TMT-LC/LC-MS/MS. **b,** Principal component analysis (PCA) of top 5% of most variant phosphopeptides. **c,** Heatmap of the changed phosphosites of LRRK2, RAB12, and OXR1. The LRRK2 data are not available (NA) in the KO mice. The phosphosites were selected based on an FDR of less than 0.05 and were subjected to manual examination. Additionally, the intensities of these phosphosites were normalized to the protein abundances from the whole proteome analysis. **d,** Immunoblot analysis of striatal tissues from WT, WTtg, GStg, and *Lrrk2* KO mice. The levels of phospho-RAB12 (pRAB12), total RAB12, phospho- LRRK2SS935 (LRRK2S p-S935), and total LRRK2 were detected. Each dot in the graphs represents an individual mouse. Each dot (**b, d**) or column (**c**) in the graphs represents an individual mouse. One-way ANOVA followed by Turkey’s HSD post hoc tests, \*\*\*\**P*<0.0001; error bars = SEM; scale bars are indicated. (**a**) was generated by BioRender (https://www.biorender.com/).

**Fig. S2.**
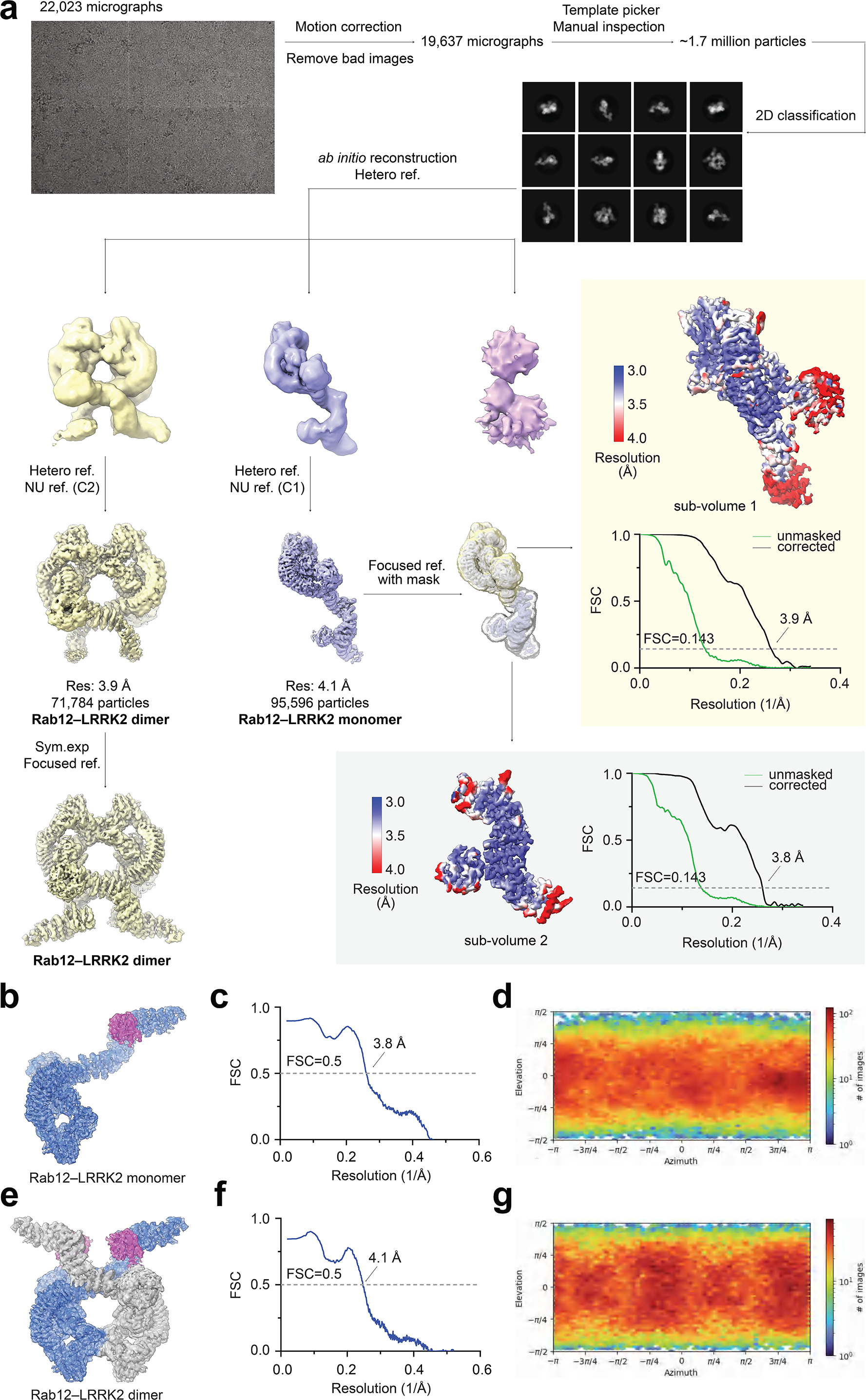
**Cryo-EM analysis of the RAB12–LRRK2 complex. a**, A simplified flow chart of cryo-EM data processing. Focused refinement was performed for two sub-volumes. Sub-volume 1 contains the LRRK2 LRR-ROC-COR-KIN-WD40 domains, and sub-volume 2 contains the LRRK2 ARM domain in complex with RAB12. Local resolutions are calculated and shown. **b**,**e**, Composed RAB12–LRRK2 maps in the LRRK2 monomer (**b**) and dimer (**e**) states with atomic model docked in. A composed map of the RAB12–LRRK2 complex in the LRRK2 monomer state was calculated in UCSF Chimera by combining sub-volumes resulted from focused refinement after aligning with the full complex map in (**a**). **c**,**f**, The FSC curve between the RAB12–LRRK2 model and composite map in the LRRK2 monomer (**c**) and dimer (**f**) states. **d**,**g**, Angular distribution calculated in cryoSPARC for particle projections (Heat map shows number of particles for each angle) of the RAB12–LRRK2 complex in the LRRK2 monomer (**d**) and dimer (**g**) states.

**Fig. S3.**
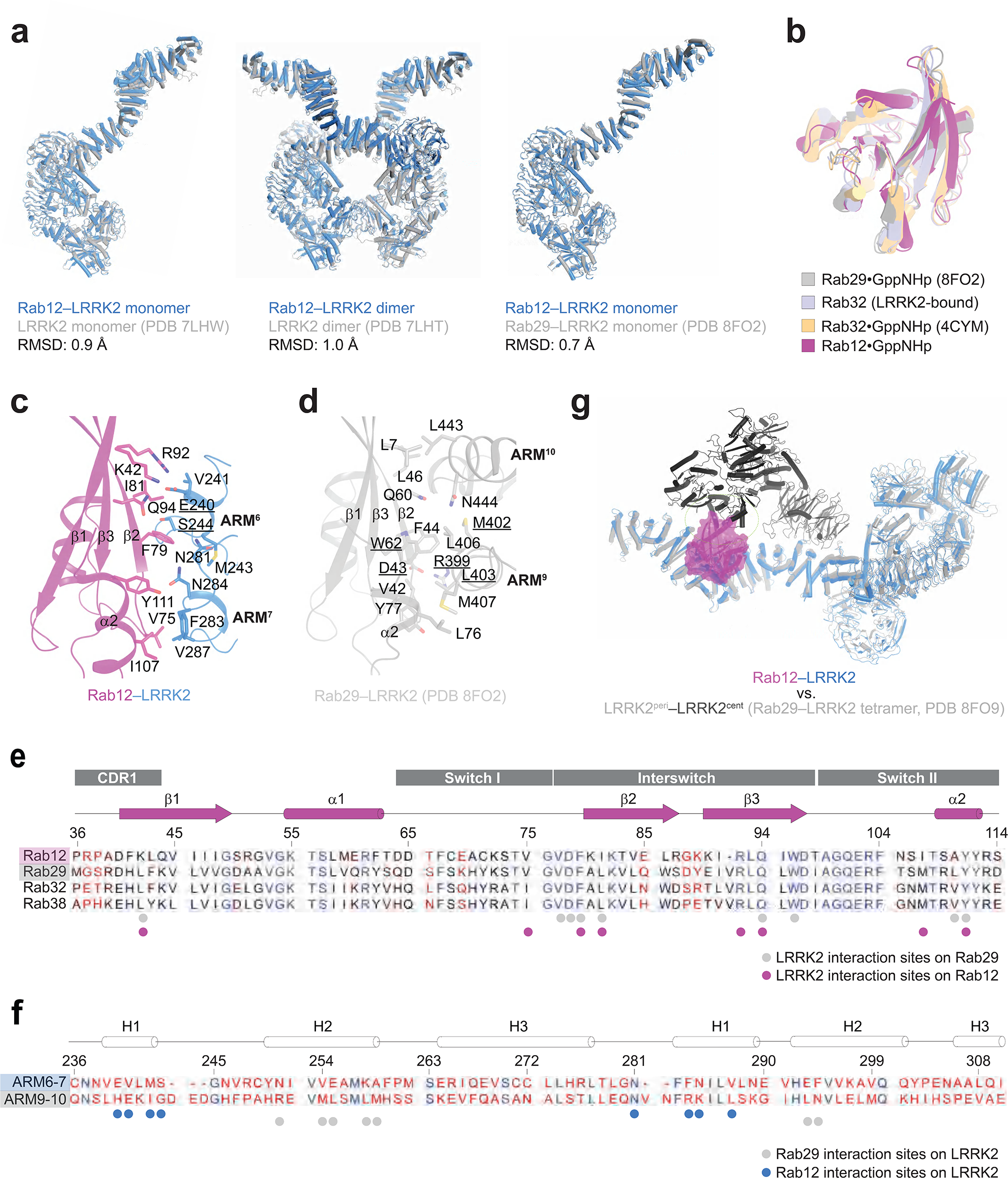
**Structural features of the RAB12–LRRK2 complex. a**, LRRK2 comparison between RAB12–LRRK2, LRRK2 alone and RAB29–LRRK2 complex structures. Left: RAB12–LRRK2 monomer and LRRK2 monomer (PDB 7LHW). Middle: RAB12–LRRK2 dimer and LRRK2 dimer (PDB 7LHT). Right: RAB12–LRRK2 monomer and RAB29–LRRK2 monomer (PDB 8FO2). **b,** Superposition of LRRK2-bound RAB12 with RAB29 and RAB32. **c-d**, Details of the RAB12–LRRK2 (**c**) and RAB29– LRRK2 (PDB 8FO2) (**d**) interfaces. Key residues involved in the interaction are underlined. **e**, Sequence alignment of RAB12 and RAB32 subfamily members (RAB29, RAB32, and RAB38). The residues involved in the interaction with LRRK2 are indicated with circles. The secondary structures and “switch” motifs are also labeled. **f**, Sequence alignment of the LRRK2 ARM6-7 and ARM9-10 repeats. The three helices of each ARM repeat are labeled sequentially. **g**, Superposition of the RAB12–LRRK2 monomer and RAB29–LRRK2 tetramer (PDB 8FO9). The LRRK2-bound RAB12 molecule is shown as a cartoon model in a transparent surface. The steric clash between LRRK2-bound RAB12 and LRRK2cent in the RAB29–LRRK2 tetramer is indicated by a dashed oval. For simplicity, only half of the RAB29–LRRK2 tetramer is shown.

**Fig. S4.**
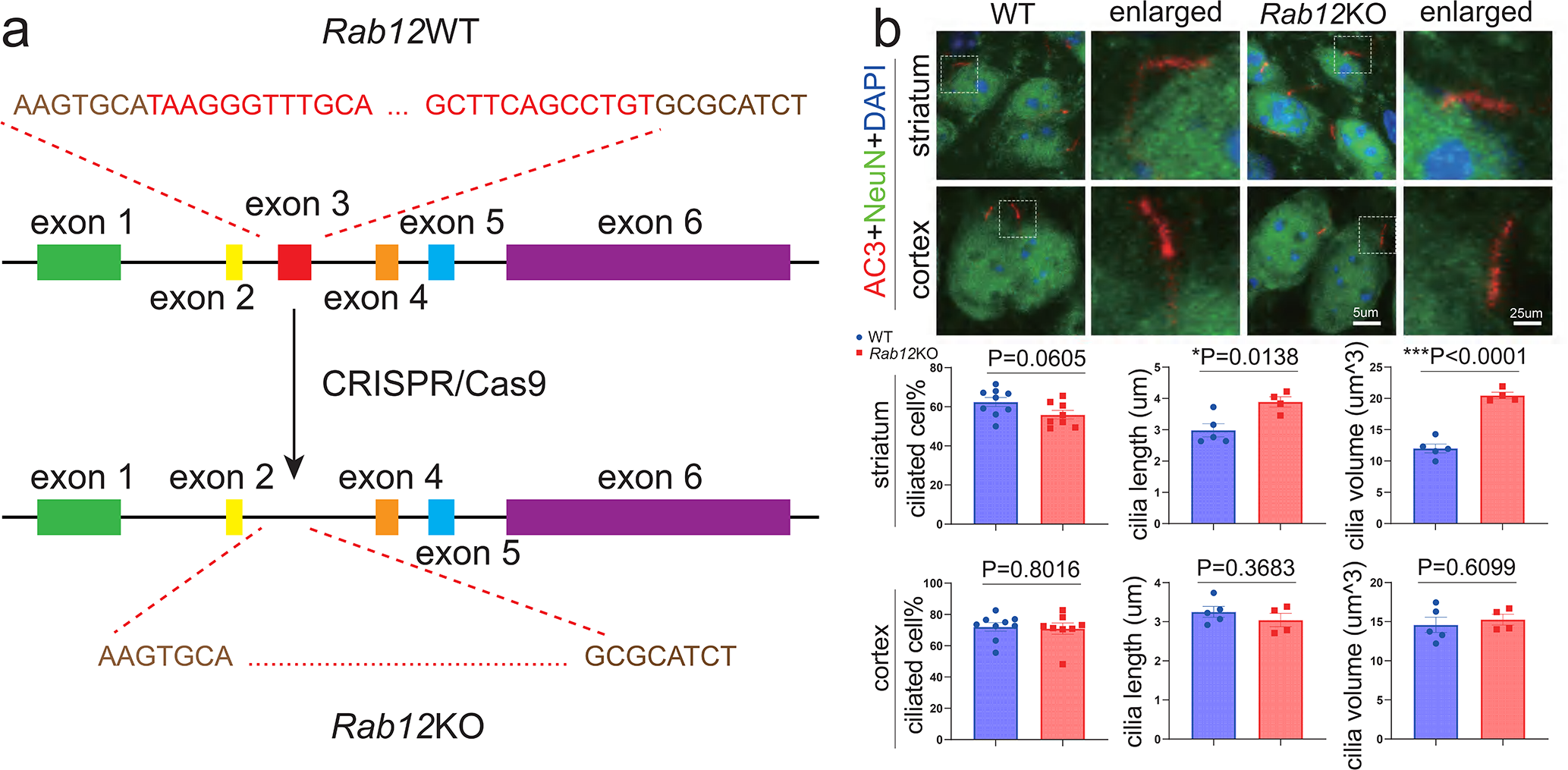
**Characterization of the neuronal ciliogenesis in the striatum and cortex of *Rab12* KO mice. a**, The schematic of the strategy for *Rab12* KO in the mouse genome. The exon 3 of *Rab12* gene was targeted and deleted by CRISPR/Cas9 technology. The sequences at the top and bottom of the *Rab12* gene schematic represent the targeted sequence in exon 3. **b**, The neuronal cilia in the striatal and frontal cortical areas of *Rab12* KO mice and the controls were detected by IF using antibodies labeling neuron-specific cilia marker AC3 and neuron marker NeuN. The boxed images were enlarged and displayed in the last image column accordingly. Each dot in the graphs represents an individual mouse. Each dot in the graphs represents an individual mouse (**b**). Unpaired two-tailed *t*-tests, \*\*\**P*<0.001; error bars = SEM; scale bars are indicated.

**Fig. S5.**
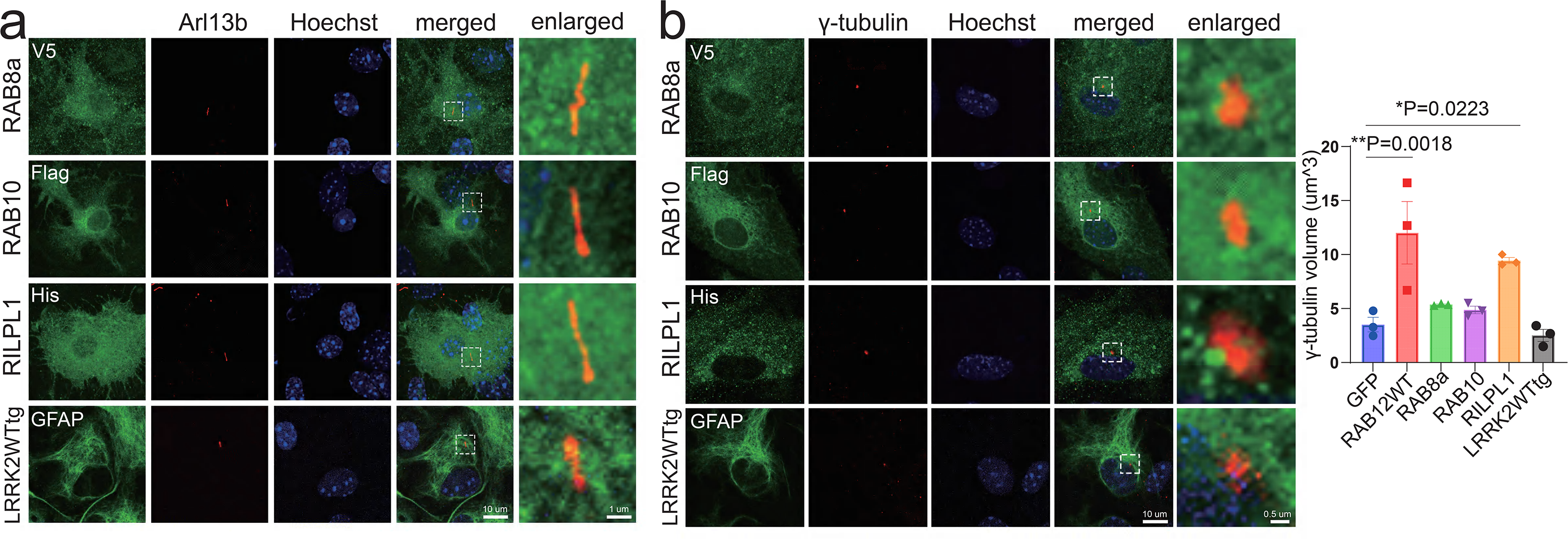
**The effects of LRRK2, RAB8a, RAB10, and RILPL1 on astrocytic cilia morphology and centrosomes homeostasis. a**-**b**, WT primary culture of astrocytes was infected with lentiviruses carrying GFAP promotors to transduce *Gfp*, HA-*Rab12*WT, V5-*Rab8a*, Flag-*Rab10*, or His-*Rilpl1*. The astrocytes from neonatal mice overexpressing *Lrrk2* WT were cultured. Arl13b (**a**) and γ-tubulin (**b**) in the astrocytes were detected by IF. Centrosome volume of the infected astrocytes in images in (**b**) was calculated. The images from astrocytes overexpressing GFP or HA-RAB12WT are not shown. The boxed regions are enlarged to display the representative cilia (**a**) and centrosomes (**b**). Each dot in the graph represents >60 cells from each mouse. One-way ANOVA followed by Turkey’s HSD post hoc tests, \**P*<0.05, \*\**P*<0.01; error bars = SEM; scale bars are indicated.

**Fig. S6.**
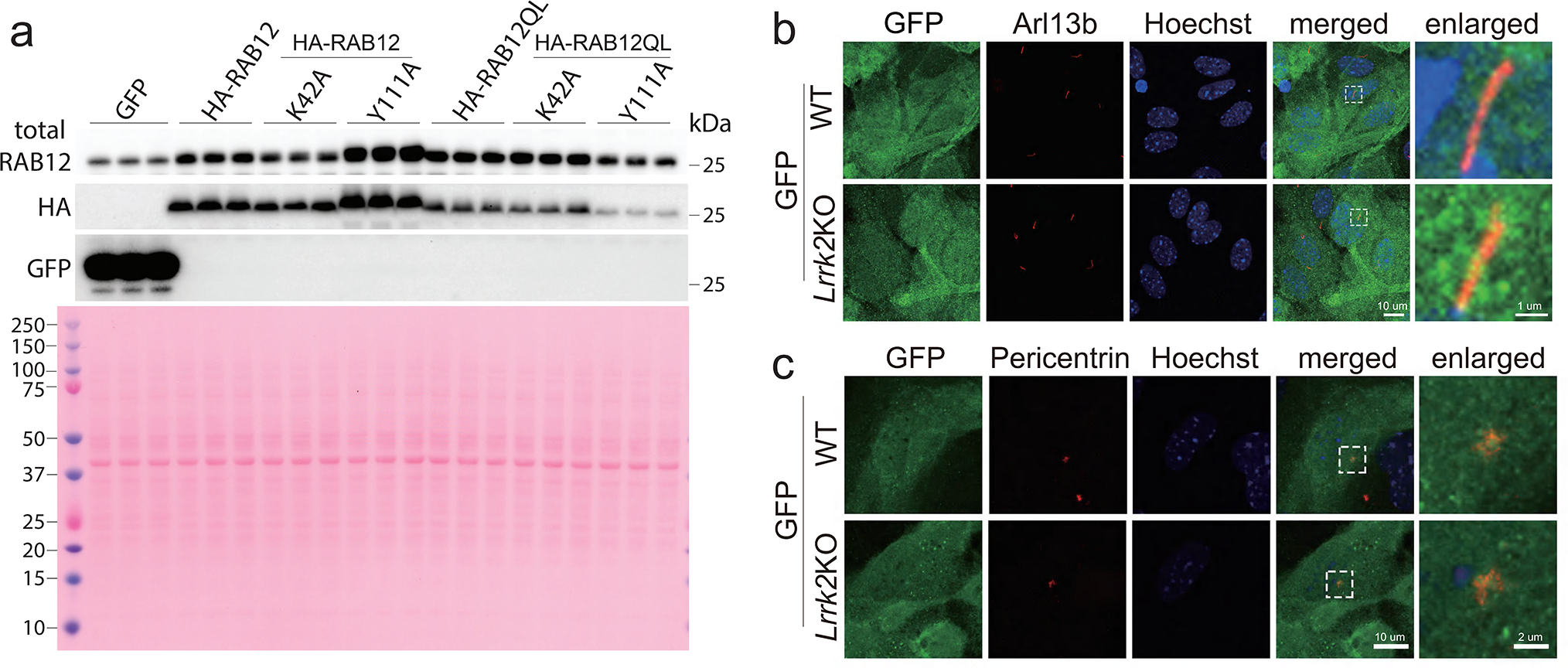
IF images of astrocytes overexpressing GFP in support of. Fig. 4d and g**-h. a,** Immunoblot analysis of primary astrocytes overexpressing GFP, HA-Rab12, HA-Rab12-K42A, HA-Rab12-Y111A, HA-Rab12QL, HA-Rab12QL-K42A, HA-Rab12QL-Y111A. The levels of total RAB12, HA, or GFP were detected. Ponceau staining was used to verify that an equal amount of protein was loaded in each lane. **b**-**c**, IF staining was applied to detect the cilia (**b**) and the centrosomes (**c**) of WT and *Lrrk2* KO primary cultured astrocytes overexpressing *Gfp*. (**b**) supports Fig. 4d, while (**c**) supports Fig. 4g**-h**. The boxed areas are enlarged. Scale bars are indicated.

**Fig. S7.**
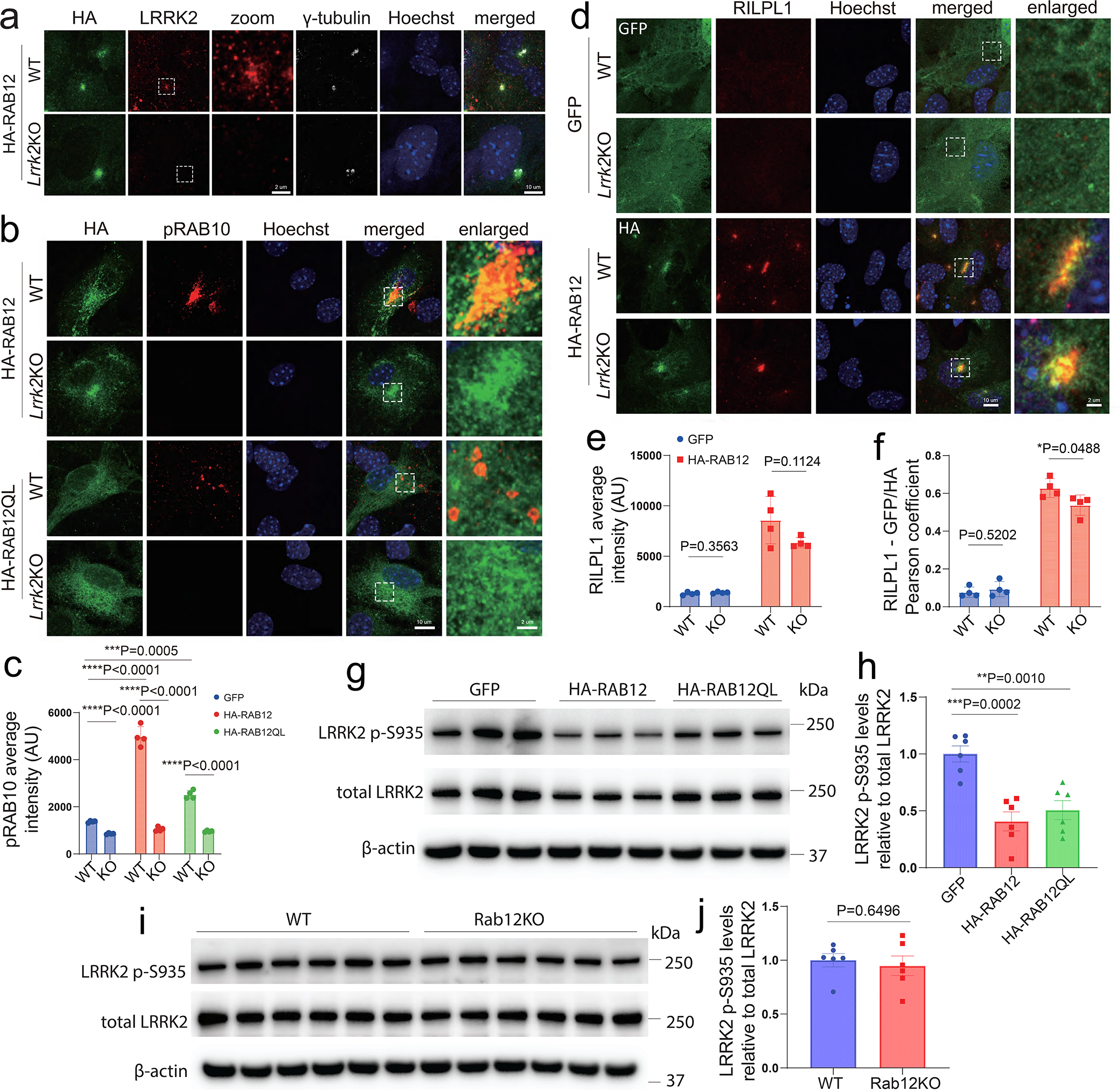
**The effect of LRRK2 on RAB12’s pRAB10/RILPL1 recruitment in astrocytes. a**, LRRK2 antibody’s specificity was validated with WT and *Lrrk2* KO astrocytes overexpressing HA-RAB12WT by IF staining. **b**-**c**, pRAB10 average intensity colocalized with GFP or HA was measured in WT and *Lrrk2* KO astrocytes overexpressing HA-RAB12WT or HA-RAB12QL by IF staining. **d**-**f**, RILPL1 average intensity colocalized with GFP or HA and RILPL1-GFP/HA Pearson coefficient were detected in WT and *Lrrk2* KO astrocytes overexpressing GFP or HA-RAB12WT. Magnification of the boxed areas in (**a**, **b**, **d**) was presented in the “enlarged” in each panel. **g-h**, Immunoblot analysis of primary astrocytes overexpressing GFP, HA-Rab12, and HA-Rab12QL. The levels of LRRK2 p-S935, and total LRRK2 were detected. Each dot in the graph represents >60 cells from each mouse (**c, e-f**) and cells from an individual mouse (**h, j**). Statistical analysis was performed using unpaired two-tailed *t*-tests (**c, e-f**) and One-way ANOVA followed by Turkey’s HSD post hoc tests (**h, j**). \**P*<0.05, \*\*\**P*<0.001, \*\*\*\**P*<0.0001; error bars = SEM; scale bars are indicated.

